# Material-mediated histogenesis using mechano-chemically microstructured cell niches

**DOI:** 10.1101/2021.02.10.430691

**Authors:** Peter L. H. Newman, Queenie Yip, Pierre Osteil, Tim A. Anderson, Jane Q. J. Sun, Daryan Kempe, Maté Biro, Jae-Won Shin, Patrick P.L. Tam, Hala Zreiqat

## Abstract

Stem-cell derived tissue models are commonly cultured under globally-delivered stimuli that trigger histogenesis via self-organizing activity. However, the culture of such tissue models is prone to stochastic behavior, limiting the reproducibility of cellular composition and resulting in non-physiological architectures. To overcome these shortcomings, we developed a method for printing cell niche microenvironments with microstructured cues that mediate local histogenic processes, including mechanosensing and differentiation of selected cell types. Microstructured cues include independently tunable mechano-chemical properties, with conjugated peptides, proteins, and morphogens across a range of Young’s moduli. By rationally designing niches, we mediate the structure of tissues derived from stem-cell-progenitor sources, including a bone-fat assembly from stromal mesenchyme, and embryonic tissues derived from hiPSC. We show that microstructured cues can recapitulate mechano-chemical signals resembling early embryonic histogenesis. This outcome includes a role for niche mechanics in human embryonic organization, where soft niche mechanics bias markers of mesendodermal differentiation and epithelial-to-mesenchymal-transition (EMT), as well as a demonstration of a material-mediated morphogen signaling centers able to induce foci of mesenchymal and EMT differentiation. Thus, microstructured materials can mediate local histogenic processes to enhance the structure and composition of tissue models.

---

Histogenesis proceeds by integrating mechanical, chemical, and topological information. Such ‘mechano-chemical’ information directs the signals through which diverse cell types, tissues, and organs emerge^1,2^. While leveraging developmental processes can yield tissues and organ models with complex structure^3–6^, current approaches rely on homogeneous signaling cues, such as globally administered morphogens and biomaterials with controllable bulk properties. Such global stimuli may preclude the generation of triploblastic tissues with correspondingly diverse structures and cellular functions. Further, ‘self-organizing’ cellular feedbacks initiated from global stimuli are prone to stochastic behaviors, generating tissues and organ models that are structurally irreproducible with malformed architectures. Critically, such architectures do not reproduce the structure-function relationships present in tissues and organs, and are thereby non-permissive to integrative cell, tissue, and organ functions. One method to overcome this limitation is to support cell and tissue differentiation with heterogeneous microenvironments that mimic the cell niche’s mechanical^7–9^ and chemical^10–14^ complexities. Here we developed technologies that can recapitulate complex cell niche environments and, in turn, specify the mechano-chemical signals to enhance the structure of multilineage tissue constructs.

One promising technology to generate complex niche environments is 3D printing. Standard printing methods involve the extrusion of a biopolymer ink or the direct extrusion of cells. Extrusion printers can produce materials with discrete properties through the sequential extrusion of different materials, either from separate print cartridges, each loaded with different bioinks^15–19^, or through mixing solutions before their extrusion through a single nozzle^20,21^. Demonstrated structured properties include changes to embedded cell types^18,22^, mechanical properties^17,21^, and biochemicals^23^. Structured materials are analogous to a pixelated digital image, where finite-area elements of specific local properties – e.g., in the case of an image: specific colors – are combined to achieve an emergent function. While the pixel organization of digital images reconstructs visual information, 3D printed multi-or-structured materials can function as a device. Structured materials fabricated with extrusion printing are limited in the number of different material types by the number of print cartridges or their ability to efficiently switch and mix solutions within a single print cartridge. Accordingly, structured cell systems fabricated from extrusion printing have been limited to no more than three cell inks^19,24^, and have not been demonstrated with comparable feature sizes to those possible through alternative printing methods, including photolithography^25^. These limitations make the extrusion approach problematic for fabricating biomaterials with microstructured properties. As an alternative approach to indirectly structuring tissues with environmental cues, organoid models have been generated using direct extrusion printing of hiPSCs^22,26,27^. hiPSCs can be directly extruded into defined architectures and subsequently differentiated on the provision of external media-derived cues or intercellular feedback from concurrently extruded differentiated cells. While this approach permits the generating macroscale organoids with improved reproducibility, these models still rely on self-organizing processes, limiting their size and tissue complexity constraining the acquisition of higher-level cellular functions.

Photolithography is an alternative 3D printing method that uses light to selectively polymerize a material from a photoresist. This method offers technical solutions to some obstacles faced when extrusion-printing complex materials. For example, photolithographic printing methods have generated materials with nanoscale features^28^, a feat yet to be achieved using extrusion-printing. Early work developing heterogeneous biomaterials with chemically discrete properties used photolithography to pattern regions of small bioactive molecules and peptides^29–31^, with more recent methods capable of the discrete spatiotemporal micro-structuring of sensitive and complex biomacromolecules^32^, and discrete mechanical microproperties^33^. Photolithographic methods print materials with heterogeneous properties by changing the composition of a photoresist during printing by serial injection of photoresists with different composition^34^. These photoresists flow through the polymerization volume hence the name flow lithography (FL), a method that has been widely used in microfluidics to fabricate microparticles^35^. FL overcomes the technical hurdles of extrusion methods by using separate subsystems for solution injection, mixing, and placement/polymerization, whereby materials with structured properties can be fabricated with a resolution primarily limited by the focus of light, the so-called ‘spot-size’. While previous works using photolithography have generated niche environments that locally regulate cell attachment^25,31^ and proliferation^32^, their capacity to define tissue structure by guiding histogenic processes, including mechanosensing and cell differentiation, remains unexplored and represents a critical development towards generating synthetic tissues, as outlined in numerous recent reviews^36–42^.

The present study establishes materials-based strategies for mediating histogenesis using mechano-chemically microstructured niche environments. We demonstrate niche material-mediated cellular-scale mechanosensing and the differentiation of selective cell types. Niches with specific microstructured properties can support the generation of stem-cell-derived tissue constructs such as a bone-fat-assembly from stromal progenitors. While exploring mechano-chemically structured niches with hiPSCs, we revealed a potential role of niche-mechanics in directing germ layer histogenesis where the embryonic tissues organize into polarized micropatterns of stem cell derivatives reminiscent of germ layer differentiation during early embryogenesis, as well as the first demonstration of a material-mediated morphogen signaling center, showing the induction of foci of mesenchymal and EMT differentiation.

## Results

### Printing microstructured niches with mechano-chemical flow lithography

Cell niche environments were printed by flowing photoresists of variable biochemical and polymeric composition through a chamber during printing (**Fig. 1, S1-3**). The photoresist is selectively polymerized using a 405 nm laser, with changes to the laser’s focus controlling the resolution and size of printed structures (**Fig. 1b-1, S4**). The photoresist is composed of a bioinert hydrogel monomer (polyethylene glycol diacrylate), a photoinitiator (lithium phenyl-2,4,6-trimethylbenzoylphosphinate or LAP^43^), and biochemicals, which can include any peptide, protein, or morphogen with a thiol-functional-group available from cysteine peptide moieties (**Fig. 1c-e, S5**). Following laser-induced photocleavage of the photoinitiator, the hydrogel monomer polymerizes alongside a thiol-ene bioconjugation reaction that can covalently crosslink biochemicals to the otherwise bioinert material, specifying bioactivity. Relatively stiff (**Fig. 1b-3, f)** or soft (**Fig. 1b-5, g**) Young’s moduli are achieved by respectively increasing or decreasing the monomer and photoinitiator concentration (**Fig. 1f-g, 2d-e**). Coordinating the above, mechano-chemical flow lithography (MCFL) can print 3D architectures that support cell attachment and growth (**Fig. 1h, i**).

**Fig. 1.**
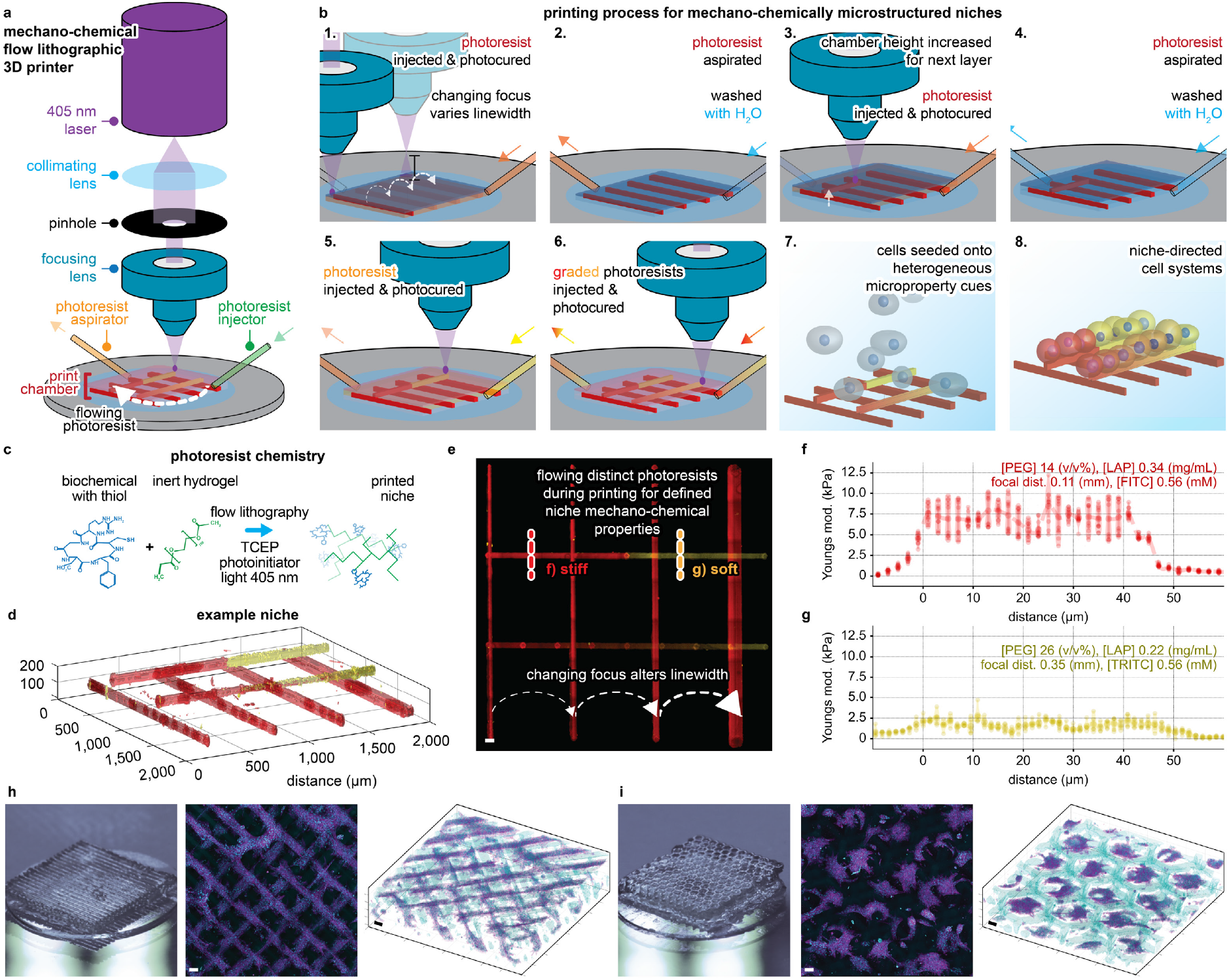
Mechano-chemical flow lithographic (MCFL) printing of microstructured niches. **a**, Components of the MCFL 3D printer. **b**, Stepwise fabrication processes to print synthetic cell niche environments. **c**, Photoresist chemistry used. **d**, Confocal image of an example niche with microstructured properties, including changes to linewidth, mechanics (Young’s modulus), and chemical microproperties (concentration of the fluorophores FITC and TRITC). **e**, Maximum intensity projection of the 3D confocal data, scale bar 50 µm. Dotted red and orange lines annotate the profile of force spectroscopy in **f**,**g**. Young’s modulus across filaments. Fabrication variables are shown at the top of the graphs for physiologically **f** stiff (7.5 kPa, red, TRITC) and **g** soft (2.5 kPa, orange, FITC) segments. **h**,**i**, Actin (magenta) and hydrogel (cyan) stained ADSCs (primary human adipose-derived stromal cells) cultured over 3D niche with **h**, ‘stacked-logs’ or **i**, ‘offset-honeycomb-layers’ architecture. Macro lens photography (left) is shown together with MIP (middle) and 3D confocal renders (right), scale bars 200 µm.

To print materials with independently tunable mechano-chemical properties, we developed a model that relates the MCFL fabrication variables to the printed linewidth, Young’s modulus, and the concentration of conjugated thiol-ene biochemical of the printed hydrogels. We characterized six variables affecting material properties, including three variables controlled by the printer (laser scan velocity, focus, and laser power) and three photoresist variables (the concentration of photoinitiator, monomer, and bioconjugate Biotin-PEG-SH, a model biochemical with free thiol group) (**Fig. 2a-f**). Decreased light exposure (**Fig. 2a, b**), photoinitiator (**Fig. 2d**), and monomer concentrations (**Fig. 2e**) decreased both Young’s modulus and the linewidth of prints, consistent with a lowered rate of polymerization due to the reduced light absorption and consequentially lower photoinitiator cleavage and monomer conversion (see notes on photopolymerization in the supplementary information, **Fig. S6-7**). The diameter of the conic angle of laser transmittance linearly correlated to changes in linewidth (**Fig. 2c**). We further characterized the effect of changing Biotin-PEG-SH concentration on material Young’s modulus and linewidth. We found that Biotin-PEG-SH inclusions up to 8 mM did not significantly alter Young’s modulus or linewidth (**Fig. 2f**).

**Fig. 2.**
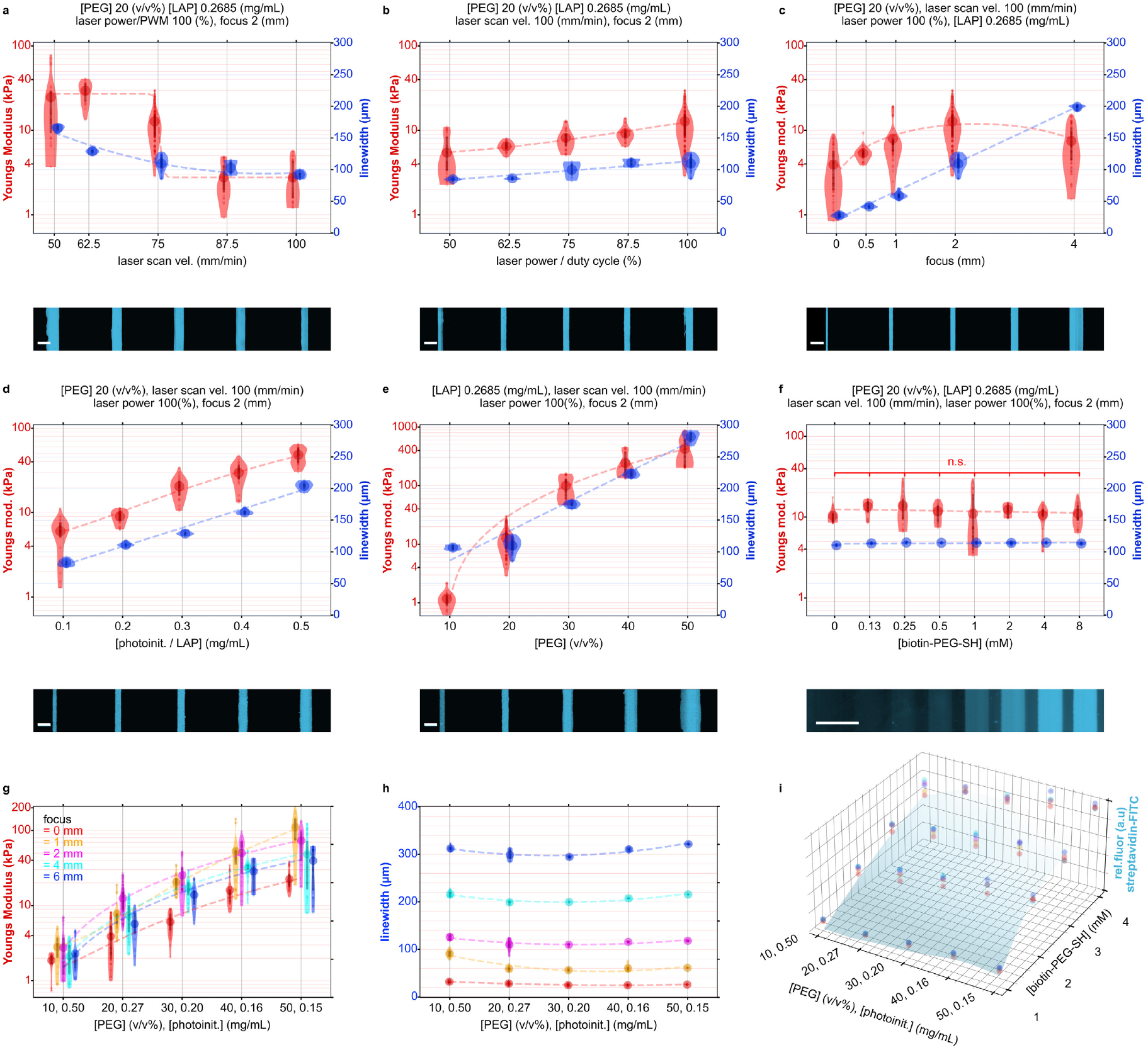
Model for printing mechano-chemically microstructured properties. **a-f**, Microstructured niche properties were printed using a model that relates printer and photoresist variables to the Young’s modulus (left axis, red) and linewidth (right axis, blue). Independent variables are indicated at the subfigure base above microscopy of printed filaments with streptavidin-FITC conjugate. Gamma correction is applied to subset **f** for improved visibility. Fabrication variables that remain constant are enumerated at the top of each subset. Scale bars, 200 µm. **g-i**, The optimized variable state-space. Discrete material properties can be interpolated from the optimized state-space to generate niches with microstructured properties. **g**, The relationship between the unified photoinitiator-monomer concentration (horizontal axis) and focus (shown in different colors) on Young’s modulus and **h**, linewidth. **i**, The effect of photoinitiator-monomer (horizontal axis), Biotin-PEG-SH concentration (into-page), and focus (shown in different colors) on the bound thiol-ene conjugate. n.s. – p > 0.05 by one-way ANOVA with Tukey post hoc tests. Violin plots show mean with 1^st^/3^rd^ quartile lines. Splines-of-best-fit are plotted to highlight trends. Throughout figure **g**, data points are offset on the horizontal axis to minimize overlap of the concurrently shown dependent variable Young’s modulus.

A variable state-space that simplifies the number of independent printing variables while still exhibiting physiologically relevant material properties was characterized (**Fig. 2g–i**). This simplification was achieved by fixing the laser scan velocity to 100 mm min^−1^ and the laser power to 100%, as well as by unifying the two variables of photoinitiator and monomer concentration to a single variable by changing the concentration of one as a function of the other (Methods, Equation 2). These simplifications left a state-space with the three independent variables: focus, bioconjugate concentration, and the combined concentration of photoinitiator–monomer. Niches with microstructured properties could then be printed by interpolating the relevant fabrication variables from the simplified state-space and discretizing the printing processes (**Fig. 2g-i**, see Methods for additional details). We demonstrated this approach by printing filaments with all permutations of either increasing or decreasing Young’s moduli, linewidth, and relative bioconjugate fluorescence (**Fig. S8**). This method allowed the characterization of a state-space and subsequent interpolation of materials properties for Young’s moduli between 2-20 kPa, biochemical additives to 4 mM, and linewidths from 40 to 300 µm (**Fig. 2g-i**).

### Niche-directed cell attachment, spreading, and mechanosensing

We explored if microstructured niche properties could direct cellular-scale attachment and mechanosensing changes. To test this, we printed niches with a physiologically moderate Young’s modulus of 8 kPa^44^ in parallel filaments between 50–250 µm wide (**Fig. 3a, Table S2**). On culturing multipotent human adipose-derived stromal cells (hADSCs) for 72h, cells selectively attached to regions printed with the cell attachment peptide RGD (cyclo(Arg-Gly-Asp-d-Phe-Cys)), displaying increasing spreading over filaments of larger linewidth. We next explored if microstructured RGD concentrations could regulate cell attachment and spreading in a dose-dependent fashion. Niches structured with six regions of different RGD concentrations between 0–8 mM were printed at a moderate Young’s modulus (8 kPa) and a fixed linewidth (250 µm) (**Table S2**). Our results showed that the extent of cell attachment and spreading correlated with the RGD concentration (**Fig. 3b**), demonstrating that microstructured niche chemistry can define local changes to cell attachment and spreading.

**Fig. 3.**
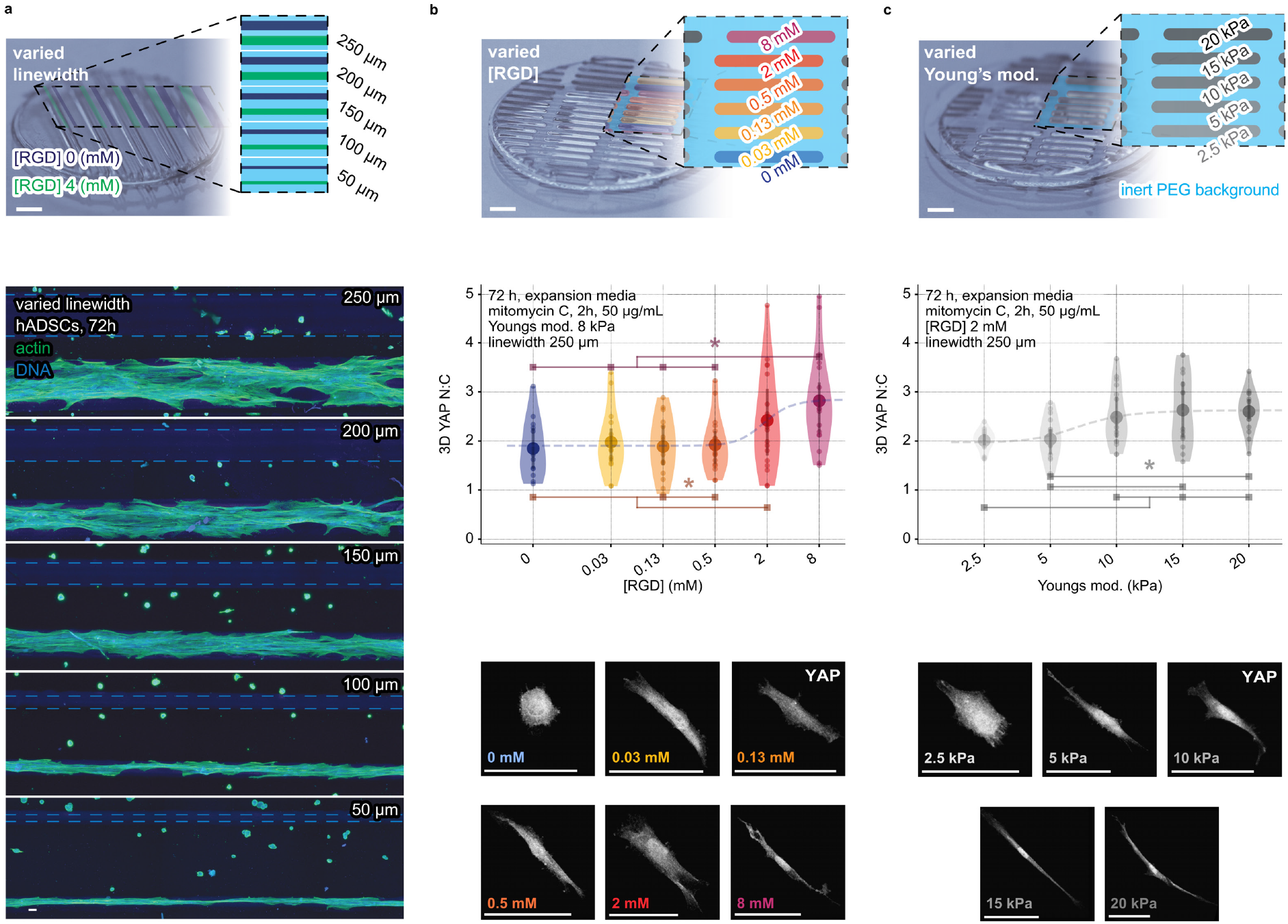
Niche-directed cell attachment, spreading, and mechanosensing. The effect of microstructured niche properties, including the concentration of the peptide cell-attachment ligand RGD and Young’s modulus, is shown to regulate cell attachment, spreading, and mechanosensing. **a**, Printed filament-pairs with 4 or 0 mM RGD, from 50–250 µm, with a fixed Young’s modulus of 8 kPa. Representative confocal MIPs show actin (phalloidin, green) and nucleus (DAPI, blue). Gamma-correction and dotted lines help improve the visibility of the zero RGD concentration dark blue filaments where no cells attach. Niche arrays with microstructured **b**, chemical (RGD) and **c**, mechanical (Young’s modulus) properties are shown alongside the quantification and representative confocal MIPs of the nuclear to cytoplasmic ratio (N:C) of the biomolecular mechanostat YAP in hADSCs at 72 h as imaged on their respective microproperty regions. Violin plots show mean with 1^st^/3^rd^ quartile lines. Sigmoidal best-fit lines are plotted alongside results to highlight trends. Scale bars, 50 µm, except macro lens images of MCFL arrays at figure top = 1 mm. Schematic overlays of material properties introduce color coding for ease of reading. *denotes p < 0.05 by one-way ANOVA with Tukey post hoc tests.

We next postulated that microstructured niches could spatially direct more complex cell functions, including cellular-scale changes to mechanosensing. We sought to replicate seminal works exploring mechanosensing^45^, in the context of our method, using local microstructured mechanical cues. To test this, we measured the localization of YAP/TAZ, a protein that acts as a mechanostat when comparing the relative distribution of YAP/TAZ in the nucleus (N) and the cytoplasm (C), or the YAP N:C ratio, wherein a high N:C ratio indicates a mechanically active cell with relatively high intracellular force^45^. As previously used, we printed niches with six chemically microstructured RGD concentrations, as well as niches with mechanical microstructure, with five regions of differing Young’s moduli between 2.5-20 kPa, corresponding to a range from physiologically soft-to-stiff (**Fig. 3c, Table S2**). hADSCs cultured on these niches displayed changes to YAP N:C ratio in a sigmoidal response to changes in the microstructured concentration, revealing that local cell mechanosensing could be regulated through the microstructure of the underlying chemical (RGD, **Fig. 3b, S9**) and mechanical (Young’s moduli, **Fig. 3c**) properties^45,46^. The lower threshold of YAP N:C correlated to a low concentration of RGD and soft Young’s modulus, while for high concentrations of RGD and stiff Young’s modulus, an upper threshold of YAP N:C was observed. This is the first demonstration of a method to define niches with complex mechano-chemical microproperties that elicit a local mechanosensing response via HIPPO signaling^47^, a critical regulator of histogenesis, morphogenesis, and organ growth.

### Niche-directed mechano-chemical assembly of a bone-fat construct

Since biological systems exhibit structure-function relationships, methods to better recapitulate physiological tissue structures *in vitro* may enable improvements to their function. We therefore explored the potential for MCFL to spatially pattern multilineage functions. Given the upstream role of cellular mechanosensing in cell fate decisions^7,45,48^, we sought to replicate seminal works regulating the multilineage functions of stromal cells, in the context of our method – using local microstructured mechano-chemical cues. Harnessing the ability for multipotent stromal cells to exhibit osteogenic and adipogenic activity^7,45^, we optimized local material properties to support concurrent functions in a structure resembling an osteon. Osteogenesis was assessed by measuring the N:C ratio of RUNX2, an essential transcriptional regulator for the commitment of stromal cells to osteoblastic and early osteogenic lineages^49^, and by visualizing mineralized bone deposition by Alizarin Red staining. Enhanced osteogenesis was revealed by elevated RUNX2 N:C ratio and enhanced mineralization, as correlated with increasing concentrations of RGD (**Fig. 4a-b**) and stiffer Young’s moduli (**Fig. 4c-d**).

**Fig. 4.**
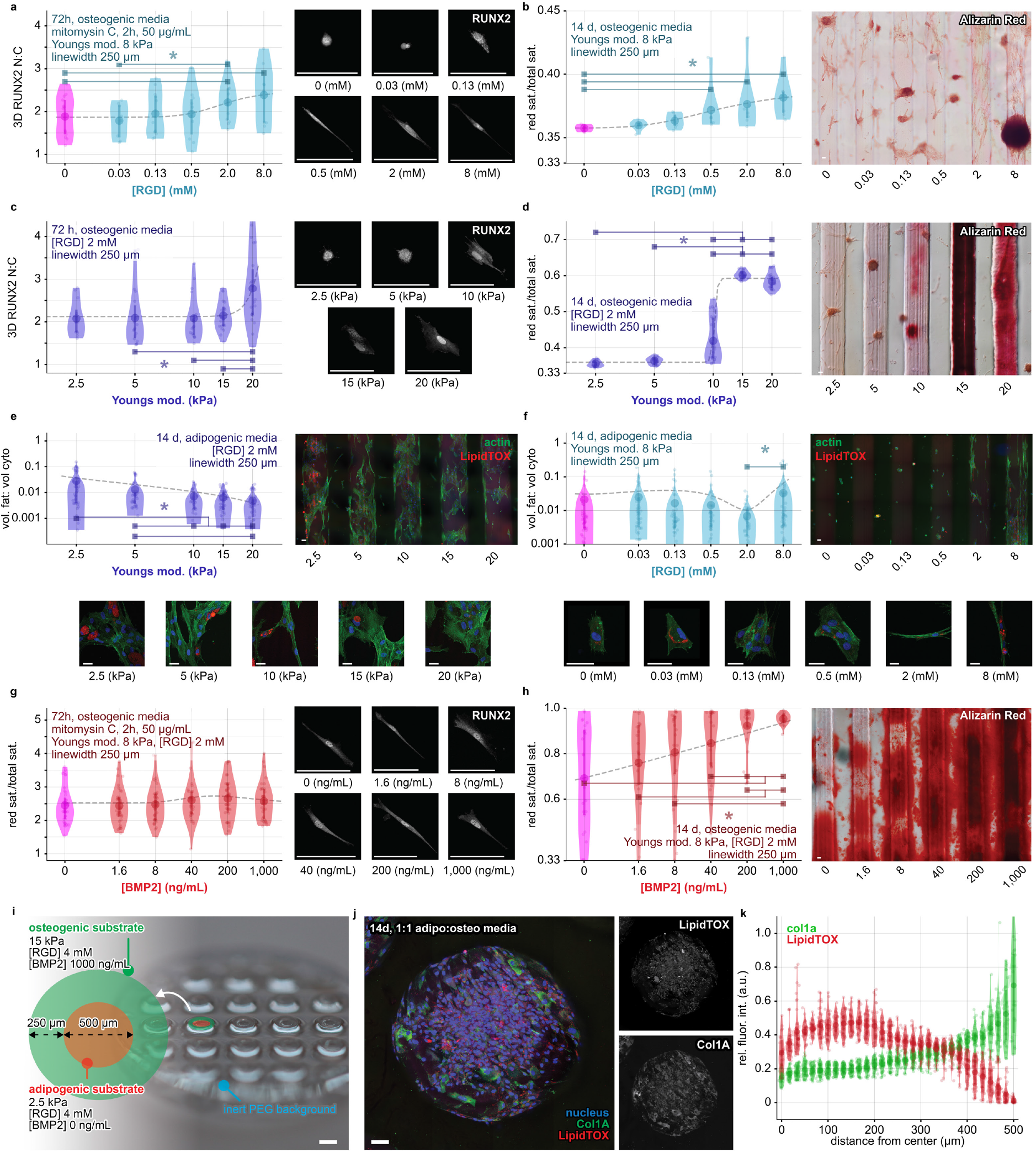
Niche-directed histogenesis of a bone-fat-assembly. Niche properties are systematically studied to engineer a bone-fat construct assembled from local material-mediated interactions. **a**,**b**, Osteogenic differentiation of hADSCs on chemically (RGD) microstructured niches. **a**, Immunofluorescent staining and quantification of RUNX2 and **b**, quantification and representative widefield images of Alizarin Red stained bone mineralization. **c**,**d**, Osteogenic differentiation over mechanically (Young’s modulus) microstructured properties, revealed by RUNX2 and Alizarin Red staining. **e**,**f** Adipogenic differentiation over mechanically (Young’s modulus) and chemically (RGD) microstructured properties, with representative MIPs and quantification of LipidTOX stained fat volume per cytoplasmic volume (vol. fat:cyto). **g**,**h**, Osteogenic differentiation on microstructured BMP2, revealed by RUNX2 and Alizarin Red stains. **i-k**, Bone-fat-assemblies that mimic the osteon with a central adipogenic region and a peripheral osteogenic region. **i**, Printed MCFL bone-fat niche array with overlaid text detailing material properties **j**, Confocal MIP image shows nuclear (blue), brightfield (grey), Col1A (green), and LipidTOX (red) channels, with grayscale insets showing Col1A and LipidTOX channels, gamma-corrected for ease of visibility. **k**, quantification of the normalized pixel intensity for Col1A and LipidTOX data across replicates. Scale bars 100 µm, except d = 1 mm. Violin plots show mean with 1^st^/3^rd^ quartile lines. Splines-of-best-fit-lines are plotted alongside results to highlight trends. Scale bars 100 µm, except for i = 1 mm. *denotes p < 0.05 by one-way ANOVA with Tukey post hoc tests.

Adipogenesis, assessed as the volumetric ratio of fat-to-total-cytoplasmic volume (fat:cyto) (**Fig. 4e-f**), was correlated with mechanically soft (Young’s modulus) or peptide-enriched (high RGD) regions. The highest fat:cyto ratio was associated with mechanical properties with 8 mM RGD and a low Young’s modulus (with ∼7-fold increase at 2.5 kPa relative to 20 kPa). Further, we tested niches with varying concentrations of BMP2, which is known to influence osteogenesis. Cells cultured over regions microstructured with BMP2 exhibit scaling of RUNX2 N:C, with increasing RUNX2 expression for concentrations up to 200 ng/mL, though decreasing at 1000 ng/mL. Mineralization, assayed by Alizarin Red activity, was enhanced with increasing BMP2 concentrations, showing that high BMP2 concentrations accelerate osteogenesis (**Fig. 4g-h**).

We next sought to direct multilineage functions of hADSCs in a well-defined structure. To elicit concurrent osteo-and-adipogenic signaling, we printed separate niche regions with optimal mechano-chemical microproperties for both osteo-and-adipogenic differentiation (**Fig. 4a-h**). A centralized soft adipogenic region with a high concentration of RGD is surrounded peripherally by a stiffer osteogenic region with a high concentration of BMP2 and RGD (**Fig. 4i**). hADSCs seeded on the bone-fat microstructured niche and cultured in a 1:1 adipogenic:osteogenic media for 14 days displayed concurrent osteogenesis (COL 1A-expressing region, **Fig. 4j**) and adipogenesis (high lipid content region, **Fig. 4j**) over their respective microstructured domains (**Fig. 4k, S10**).

While multipotency of stromal-derived cells such as hADSCs lacks rigorous evidence^50,51^, their use herein permits contextualization and comparison with seminal works that explore the relationship between material properties, mechanobiological, and multilineage functions^7,45,48^. Our demonstration builds on these seminal works showing for the first time that locally defined mechano-chemical cues can spatially guide the concurrent adipogenic and osteogenic activity of hADSCs and potentiates the coordinated generation of biologically relevant structures, potentiating defined multilineage cell systems composed of locally divergent signaling and exhibit structure-function relationships.

### Directing embryonic histogenesis by niche mechanics

During embryogenesis, changes in the mechanics of the extracellular matrix are reputed to be critical for initiating mesodermal differentiation in zebrafish^52^, avian^53^, and murine^54^ embryos. However, modulation to the mechanics of the embryonic niche remains unexplored in the context of human embryogenesis, a phenomenon that remains prohibitive to investigate. To explore the role of soft matrix mechanics during the differentiation of human embryonic tissues, we compared three types of niche environments (i) a 1000 µm diameter Matrigel-coated-glass control niche (circular control), (ii) an MCFL printed square with uniform properties throughout (uniform niche), and (iii) an MCFL square with a 1D mechano-structured gradient in Young’s modulus (mechano-structured niche) (**Fig. 5a, S11**). Cultures were stimulated with BMP4 (50 ng/mL in Essential 8-Flex, or E8F) to a 72 h endpoint and characterized using confocal microscopy with adaption of the StarDist segmentation tool^55,56^ (**Fig. 5a-b,f, S12-19**). Circular control niches replicated previous work showing the reproducible center-to-peripheral patterning of germ-layer derivatives from hiPSCs^3^. Similarly, hiPSCs cultured on square uniform niches exhibit similar marker expression patterned along the center-to-peripheral axis of the niche (**Fig. 5f-g, i-j, S14, S17**)^3^. Patterning is consistent with previous findings^3,57,58,58^, where center-to-peripheral patterning results from subsequently graded signaling cascades along the radial dimension^58,59^. Our findings contrast with previous reports that are limited to culture on materials with uniform mechanics, and show tissue types on mechano-structured niches exhibit additional phenotypic patterning bias along the mechano-structured axis (**Fig. 5g-m, S15, S18-19**). Well-spread cells populated low Young’s modulus regions of the mechano-structured niches, with further analysis revealing the preferential differentiation of cells to mesoderm (BRA, SNAI1-expressing cells) and endoderm (SOX17, FOXA2-expressing cells). In contrast, ectodermal cells (SOX2-expressing cells) populated regions of higher Young’s modulus (**Fig. 5g-m**). These findings are consistent with small-molecule inhibition studies modulating cellular tension and subsequent mesodermal differentiation^60,61^, and are confirmed with high YAP N:C, indicative of high cellular tension, in regions of mesendodermal differentiation (**Fig. 5l-m, S19**). Violin plots directly compare the distribution and relative differences in the marker expression between uniform and mechano-structured niche regions with the mean nuclear expression of all cells over each region of the mechano-structured niches normalized by the mean marker expression in the same region of uniform niches (**Fig. 5m**). Violin plots confirm a regional bias of mesodermal and endodermal cells populating niche regions with soft mechanics and ectodermal cells over stiffer mechanics, with data at 48 h revealing a similar trend (**Fig. S15**). Mesoderm and endoderm cells were localized to regions with enhanced SNAI1 expression, suggesting that mesendoderm differentiation may be accompanied by active epithelial-to-mesenchymal-transition (EMT)^62^ (**Fig. S17-19**). Differences between uniform and mechano-structured niches highlight the impact of extracellular mechanics on human germ layer differentiation. Our findings are consistent with the notion that softened extracellular matrix may induce mesendodermal differentiation and EMT, reminiscent of that in the zebrafish^45^, avian^46^, and murine^47^ embryo. Thus, the mechanics of the human embryonic matrix may play a role during embryonic tissue polarization by inducing a bias to mesodermal differentiation and the expression of EMT transcription factor SNAI1.

**Fig. 5.**
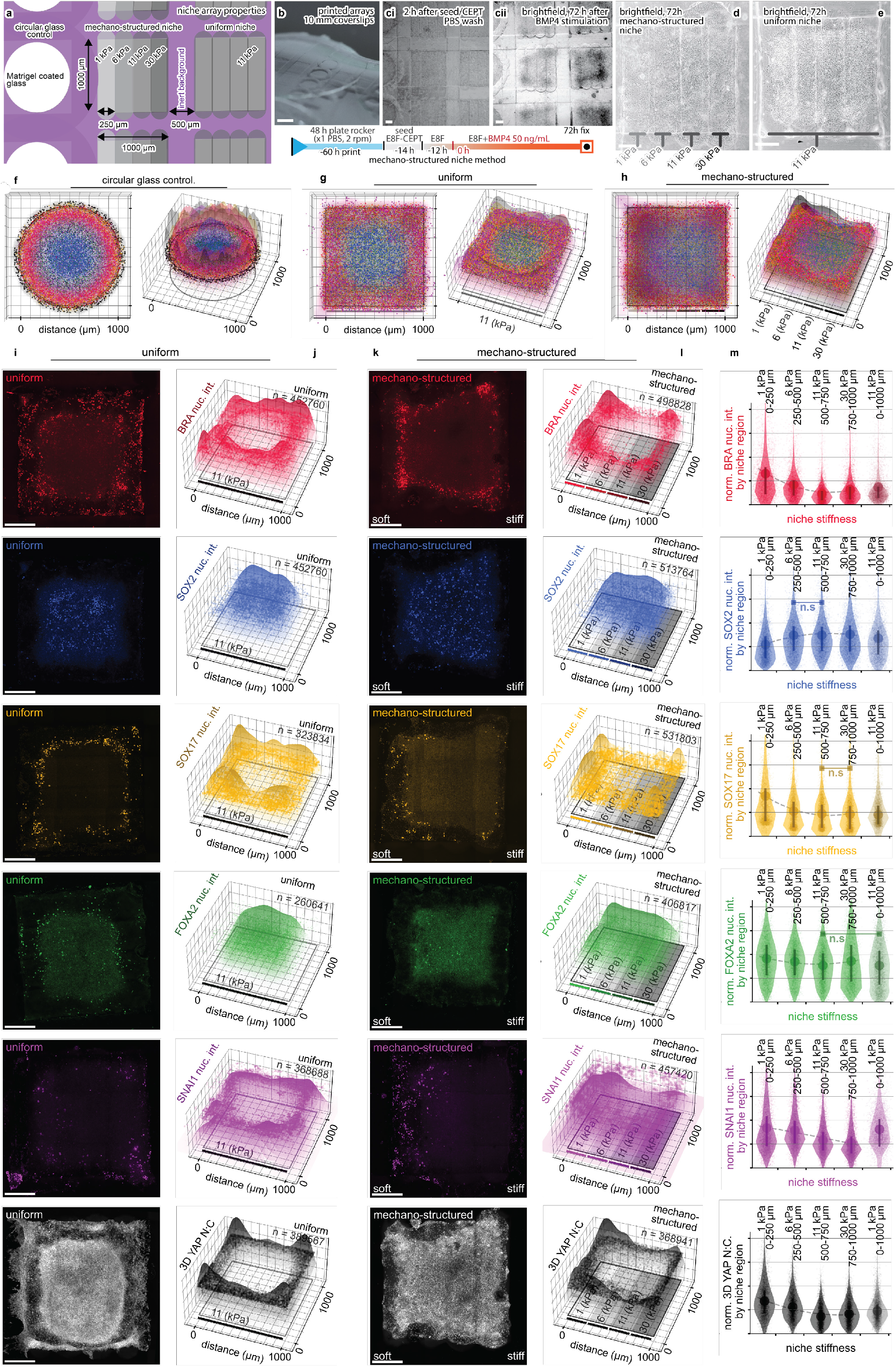
Microstructured niche mechanics recapitulate softened embryonic matrix and bias regionalized histogenesis. The response of hiPSCs cultured over circular control, uniform, and mechano-structured niches. Symmetric marker expression is observed in uniform and circular control niches, compared with marker arrangement over mechano-structured niches, which biases mesodermal (BRA-red, SNAI1-purple) and endodermal (SOX17-yellow, FOXA2-green) markers to material properties of low Young’s modulus. In contrast, ectodermal SOX2-positive cells localized to properties with higher Young’s modulus. All cell data shown at 72 h, except **c**. Orientation is consistent across the figure, with mechano-structure shown soft-to-stiff, left-to-right. Marker-color coding is consistent throughout work. **a-b**, Schematic and macro-lens photography show coverslips with printed arrays of the three niche-types with overlaid text indicating material properties. Scale bar 1 mm **c**, Brightfield imaging of arrays shown above an overview of the experimental method and timeline. **d-e**, Brightfield microscopy of mechano-structured and uniform niches, with overlaid text indicating mechanical properties. **f-h** Pooled-marker-expression of all markers shown simultaneously as 3D scatter and surface plots overlaid on a representation of the corresponding niche. Each dot represents a single cell with size, height and transparency scaled to the normalized fluorescent intensity. **Columns i**,**k** Show representative confocal MIPs images for uniform and mechano-structured niches, respectively. Scale bars 200 µm. **Column j**,**l** 3D scatter and surface plots for each marker assayed shown overlaid on a representation of the corresponding niche. **Column m**, Violin plots compare the relative distribution and mean differences of uniform and mechano-structured properties, including relative mean and 1^st^/3^rd^ quartile lines of replicate data as mapped to respective material property regions. ‘n.s.’ denotes p > 0.05 with all remaining group permutations p < 0.05 by one-way ANOVA with Tukey post hoc tests.

### Material-mediated morphogen signaling centers for directing histogenesis

Development proceeds in part by way of secreted morphogen signaling centers. For example, the vertebrate blastula can be committed to a well-developed embryo using a minimally inductive system composed of two signaling centers^53^. Therein, ventral secretion of BMP4 pairs with dorsally secreted Nodal to establish signaling cascades from which embryogenesis proceeds. We explore the functionality of MCFL niches by recapitulating the ventrally-sourced BMP4 signals in a micropatterned model of germ layer histogenesis. We printed morphogen-structured square niches composed of two discrete regions, one harboring BMP4 and the other without (**Fig. 6a, Table S2**). Unlike circular, uniform, or mechano-structured niches, morphogen-structured niches were culture with no further supplementation of factors to the culture media during subsequent differentiation (**Fig. 6b-c**). Results showed changes in the nuclear translocation (N:C ratio) of the BMP4 signal transducer pSMAD1 in response to varying concentrations of BMP4 in the morphogen-structured niche (**Fig. 6e,g, S20-21**). Cells cultured over the BMP4-structured regions exhibit an increased pSMAD1 N:C ratio indicating a positive response to the local BMP4 signal (**Fig. 6e-g, S18**). By 72 h, cells over the BMP4-structured regions formed circular foci (**Fig. 6d, S22-23**) of mesoderm (BRA, SNAI1-expressing) with a diffused scattering of endoderm cells (SOX17, FOXA2-expressing) (**Fig. 6f, h-m, S22**). Mesoderm cells packed densely within and around the fociexhibiting high SNAI1 expression (**Fig. 6h, l**), with further high YAP N:C ratio central to the foci (**Fig. 6m, S22**).

**Fig. 6.**
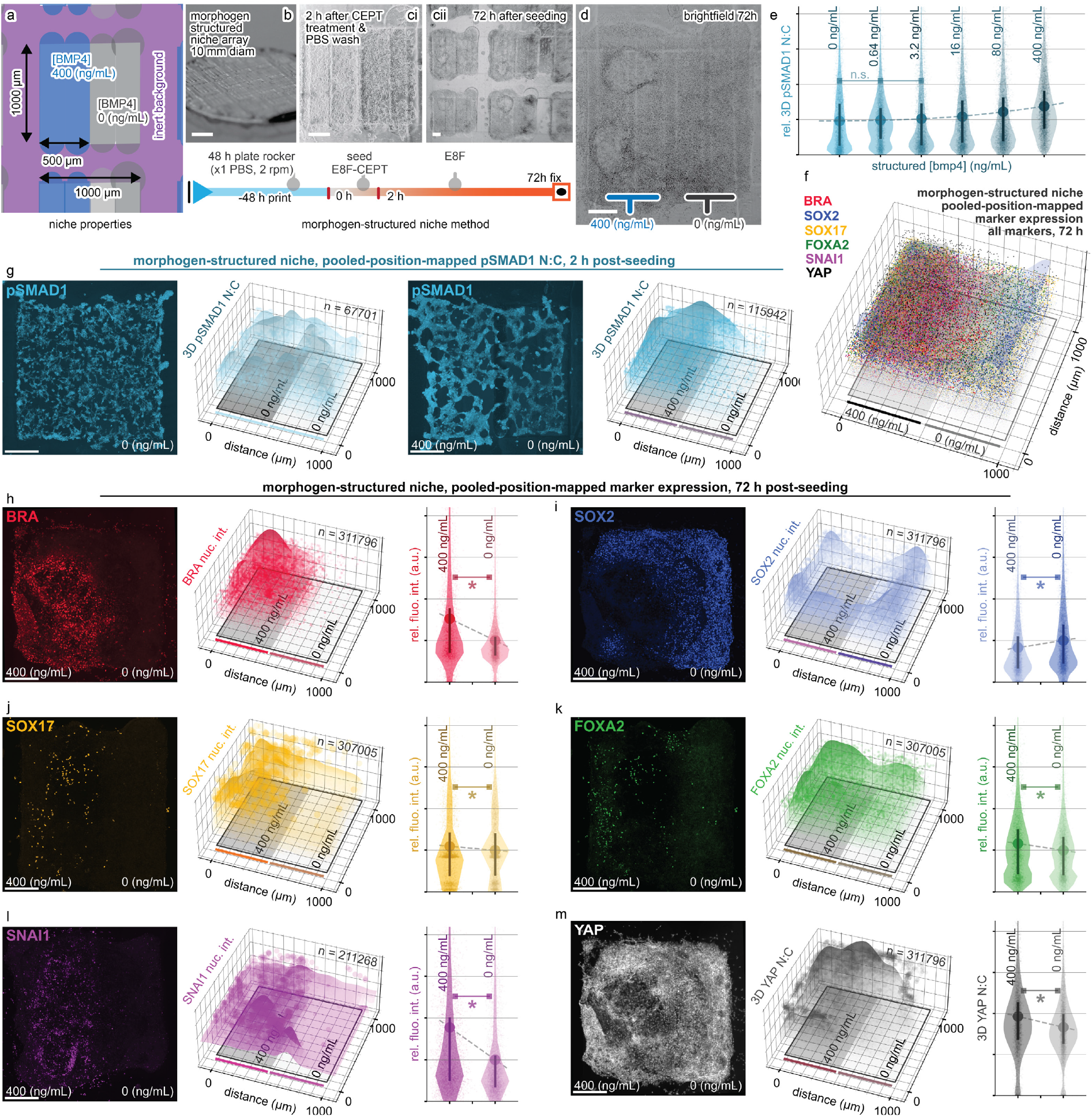
A material-mediated BMP4 signaling center directs histogenesis, inducing foci of mesenchymal and EMT markers. Histogenesis of hiPSCs cultured over morphogen-structured niches. At 72 h, circular foci appear over regions with BMP4, visible in brightfield and representative confocal images with the concurrence of phenotypes and morphology reminiscent of the a primitive-streak phenotype. Densely packed mesodermal cells (BRA-red, SNAI1-purple, YAP-gray) are surrounded by diffuse endodermal cells (SOX17-yellow, FOXA2-green), with ectodermal (SOX2-blue) cells localized to regions absent of BMP4. Orientation is consistent across the figure, with BMP4 containing regions at left and regions absent of BMP4 at right. **a**, Schematic of niche array with overlaid text indicating morphogen-structured properties. **b**, Macro lens photography of array. Scale bar 1 mm. **c**, Experimental timeline and brightfield imaging at **ci**, 2 h and **cii, d** 72 h. **e**, Violin plots comparing relative mean and distribution differences of the BMP4 signal transducer pSMAD1, in response to varying concentrations of BMP4 in the morphogen-structured niche at 2 h post-seeding. 1^st^/3^rd^ quartile lines are shown. **f**, 3D view of pooled-marker-expression maps of all markers shown simultaneously overlaid on a representation of the morphogen-structured niche. Each dot within the maps represents a single cell with size, height and transparency scaled to the normalized fluorescent intensity. **g**, Pooled-marker-maps for pSMAD1 N:C at 2h for niche absent of BMP4 (left), or a morphogen-structured niche with region of 400 and 0 ng/mL BMP4 (right). **h-m**, Representative confocal MIPs with 200 µm scale bars, 3D scatter/surface plots, and violin plots of regionalized marker expression in cultures on morphogen-structured niches at 72 h. *denotes p < 0.05, ‘n.s.’ denotes p > 0.05 by one-way ANOVA with Tukey post hoc tests.

Histogenesis induced by material-mediated BMP4 is reminiscent of poles that form during embryonic organization, including the EMT transcription factor SNAI1 and mesendoderm differentiation, as well as early time-points observed during embryonic manipulation experiments exploring the role of BMP4-Nodal signaling center pairs^63^. This demonstration shows, for the first time, material-mediated morphogen signaling centers and their application to spatially define cell differentiation, potentiating the derivation of triploblastic cell systems derived from locally divergent signaling. Morphogen-structured niches build on current stem-cell-based embryo models^3,4,36,58,60,61,64–66^, adding the capacity to explore the roles of mechano-chemical signaling and advance our understanding of how complex multimodal signals are integrated to govern development.

## Discussion

Mechano-chemically microstructured niches offer the ability to direct cell functions through defined niche-material interactions. We demonstrate that mechanically and chemically structured niche properties can elicit localized cell mechanosensing, drive differentiation of stem/progenitor cell tissue, and direct histogenesis in tissue models. This versatile method overcomes a reliance on homogenous signaling as limited by bulk material properties or traditional globally administered media-based cues^5,6^, and provides an alternatives to microfluidic systems^4,64^, microsurgical approaches^63^, and optogenetics^67^, while offering the ability to define colony shape, size, and mechanics. Therefore, MCFL microstructured niches provide an entry point for better understanding the complex multimodal mechano-chemical interactions that define tissues.

Future work using such niches may extend to multi-layer structures with increased complexity and topological constraints. Multi-layer structures can be fabricated as per **Fig. 1h-i**, using chemical photoabsorbers to simplify printing of 3D structures^68^ or multiphoton lithography. As multiphoton photolithography can achieve sub-diffraction-limited resolutions^28^, combining multiphoton methods with FL may enable printing materials with subcellular features and nanostructured properties.

Advancement to the MCFL method could be made by adopting a more specific conjugation chemistry. While our method demonstrates the function of complex biomolecules, next-generation conjugation methods may improve biochemical specificity and function. Specifically, our method is limited when conjugating relatively complex macromolecules with numerous cysteine sites. Under these conditions, thiol-ene conjugation occurs stochastically and can lead to the bioconjugate’s loss of function. This limitation has been addressed with more reproducible chemistries, including enzymatic methods^32^ and specific high affinity non-covalent binding chemistries^69^. Further, such complex chemistries address the potential for the undesired cleavage and release of thiol-ene bound growth factors that may otherwise occur. Additionally, during the generation of MCFL materials, the biological activity of a given biochemical must be considered. For example, the biological activity of some morphogens exhibit relatively short half-lives as intrinsic to their resulting morphological function. In each case, the biological activity of a given biochemical factor must be determined in coordination with the MCFL protocol, which demands niche preparation, including printing, washing, and sterilization steps.

The use of mechano-chemically microstructured niches complements current methods for generating complex stem-cell-derived tissue models. Conventionally such models use globally administered biochemical stimuli within culture media. Unfortunately, global stimuli precludes the localized differentiation of cell types, ultimately limiting tissue models from forming triploblastic tissues, along with the corresponding diversity of multicellular interactions. Our results using material-mediated stimuli in embryonic models confirm the potential to define multilineage cell systems with locally divergent signaling. The expansion of our approach in controlling localized signaling could enable the specification of tissues of divergent phenotypes and their later convergence and reintegration with complex multicellular interactions. Complex structured niches potentiate the next generation of reproducible synthetic triploblastic tissue models, where concurrent and complex histogenesis programs can be defined using niche material-mediated mechano-chemical interactions.

While it is feasible to reproducibly generate a bone-fat microtissue and polarized embryonic tissues, the continued application of these niche systems has the potential to investigate some of the biggest unanswered questions in biology, such as how complex structure and function emerge; how the shape, size, and body coordinates of an organism are determined; and how mechanical and positional morphogenetic cues works in concert with morphogens and lineage determinants.

## Methods

### Custom-built MCFL 3D printer

The MCFL printer was built from custom hardware and uses software developed by the authors. In-depth details are available in the supplementary information, listing directions for the up-to-date printer and a detailed printing method. Critical components include the high-resolution x/y and z stages (V-528.1AA /V-528.1AB and M-406 including corresponding controllers C-413 and C863 from Physik Instrumente (PI) GmbH & Co. KG), as well as 405 nm diode laser (Cobolt 06-01 Series 405 nm, fiber pigtailed, FC/APC). Various mechano-optical components were used and purchased from Thor Labs (listed **Table S1**). A copy of the printer software is included in the supplemental information, and up-to-date software can be obtained by contacting the lead authors.

### Measurement of structured niche microproperties

Niches were fabricated on acrylated coverslips using the geometries, photoresist, and printer variables as reported. The measurement of Young’s modulus was completed via force spectroscopy using a JPK. NanoWizard Sense AFM mounted on Nikon Ti microscope. The device was fitted with the SuperCut quartz cantilever holder for liquid immersion and used with Bruker MLCT pyramidal cantilevers with stiffness calibrated using the thermal noise method. For force-displacement curve generation, samples and AFM cantilever were submerged in x1 PBS. The cantilever approach velocity was fixed to 0.5 µm sec^−1^ and terminated at a threshold force of 10 nN. Measurements were taken from 3 independent experimental replicates from at least four different printed-niche-replicates in each experiment. The Young’s modulus was calculated from each force-displacement approach curve using a custom fitting program written in MATLAB, with automated contact point determination and fitting for an 18° half-angle conic section (Sneddon model, as per Bruker recommendation for MLCT pyramidal cantilevers), with sample-thicknesses bottom-effect cone-correction as per Gavara et al.^58^. Data for force spectroscopic curves of AFM tip displacement against indentation force were rejected when discontinuities in the curves were present, corresponding to samples slipping and an inaccurate indentation. For the sample shown in **Fig. 1d-g**, only two independent experimental replicates were fabricated, as this sample only served to illustrate how a mechano-chemically microstructured niche material could be fabricated with the MCFL methodology. One replicate was mounted for confocal microscopy (**Fig. 1d-e**), and the other was analyzed with force spectroscopy (**Fig. 1e-f**). Linewidth and bioconjugation were measured using confocal microscopy. Linewidth was directly measured using Fiji-ImageJ across three independent experimental replicates with quantification of the concentration of Biotin-PEG-SH was measured indirectly by measuring the relative fluorescence of bound streptavidin-FITC. Indirect measurement was used to prevent the photobleaching or free radical attack of fluorescent molecules during photopolymerization, that otherwise limited interpretation.

### Interpolation method for structured niches microproperties

Empirical data was tabulated pairing dependent and independent variables, including Young’s modulus, linewidth, and the concentration of bioconjugate (**Fig. 2g-i**), with the monomer–photoinitiator (”*PEG*/*PI*” below), focus (shown as “*Z*”) and [Biotin-PEG-SH]. Using MATLAB (2020a), we then calculated the value of the independent fabrication variables of *PEG*/*PI* and *Z* after substitution of the desired Young’s modulus (*E*_*des*._) and linewidth (*W*_*des*._) as per Equation 1 below:

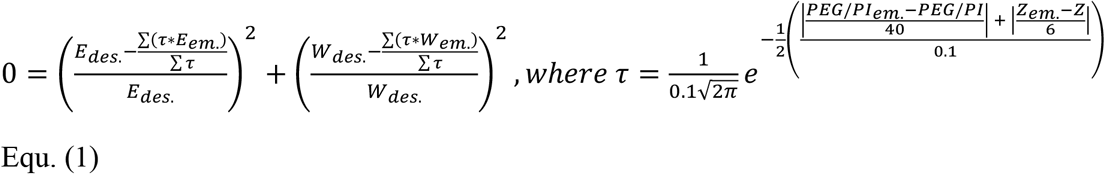

where, *PEG*/*PI*_*em*._and *Z*_*em*._ are vectors from the independent paired variables from the tabulated empirical dataset presented in **Fig. 2g,h**, and are used to calculate the vector *τ* that is substituted into Equ. 1, where a solution is obtained for the dependent paired variables *E*_*em*._, *W*_*em*._. The numbers 40 and 6 in Equ. 1 represent the normalization range for *PEG*/*PI* and *Z* respectively, over which data is interpolated. Equ. 1 is solved by finding the minimum of unconstrained multivariable functions. The relationship between *PEG*/*PI* is calculated as the positive real solution of Equ 2.

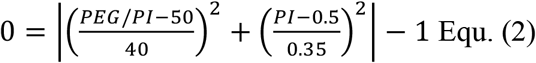

### Cell culture

hADSCs were cultured in expansion media: MesenPRO RS™ Basal Medium (Invitrogen) with the supplement of 2 mM l-glutamine and MesenPRO RS Growth Supplement (Life Technology). hADSCs at passage 3 were used for all the studies. A 10 mm coverslip with printed niche arrays was placed in a 48-well plate for culture on niches. Using a biosafety cabinet, niche arrays were washed twice with PBS before 12 mins UV-light sterilization. The medium was changed every 3 days except for studies examining YAP and RUNX2 nuclear translocation, where hADSCs were treated with mitomycin (10 µg mL^−1^) (Cayman Chemical, 11435) for 2 h, to inhibit proliferation, 24 h after seeding, after which media was replaced with expansion media. For YAP and RUNX2 translocation studies, cells were seeded at 3,000 cells/cm^2^. For differentiation studies, cells were seeded at 6,000 cells/cm^2^, osteogenic (Gibco, A1007001) and adipogenic media (Gibco, A1007201) were used as noted. Experiments using iPSCs were performed with the HPSI0314i-hoik_1 hiPSC line, obtained from the Wellcome Sanger Institute with the help of Cell Bank Australia. For routine passaging, cells were passaged with ReLeSR™ and grown on hESC qualified Geltrex coated 6-well plates. All experiments used Essential 8 Flex media (E8F), with supplements as listed, Human BMP-4 Recombinant Protein (Gibco, Catalog # PHC9534). For cell culture of iPSCs on niches, a 10 mm coverslip with printed niche arrays was placed in a 48-well plate. The niche arrays were washed for 48 h on a plate rocker in PBS before 12 mins UV-sterilization using a biosafety cabinet. hiPSC cells were dissociated with ReLeSR and pipetted into a single cell suspension with CEPT cocktail^59^ before seeding at 1 million cells per 300 µL per well, or 500k cells per 300 µL per well for morphogen-structured niches. After 2h, media was replaced with E8F, before 48h differentiation in 500 µL of E8F supplemented with 50 ng/mL BMP4 as reported (PHC9534 Gibco). Media was then replaced daily. All cells tested negative for mycoplasma contamination. For hADSCs, routine PCR assay checks (LookOut® Mycoplasma qPCR Detection Kit, MP0040A-1KT) were completed for mycoplasma contamination, for hiPSC mycoplasma was tested using the fluorescent kit, MycoAlertTM Mycoplasma Detection Kit, Catalog #: LT07-318.

### Immunofluorescence imaging and image quantification

For immunostaining, all solutions, except for those with dilute antibody and fluorophores, were syringe filtered through 0.22 µm membrane filters (Merck Millipore SLGP033RS). Cells were fixed at room temperature 4% PFA in x1 PBS buffer for 10 mins and then washed three times with PBS, followed by 12 min permeabilization at room temperature with 0.1 w v^−1^ % Triton X-100 in PBS. Samples were then incubated in a blocking buffer of 3% BSA, 3.75 mg mL^−1^ glycine, and 0.05% w v^−1^ Tween 20 in PBS for 1 h at room temperature. Primary antibodies were diluted in 1% BSA and 0.05% w v^−1^ Tween 20 in PBS and added for 2h (SOX2, 1:400, #3579, CST; SOX17, 1:400, Af1924, R&D; BRA, 1:400, AF2085, R&D; SOX17, 1:300, OTI3B10, Thermo; pSMAD1/5, 1:800, 41D10, CST; SNAIL, 1:500, AF-3639, R&D; YAP, 1:400, sc-101199, SCBT; RUNX2 AF488, 1:200, sc-390351; NANOG, 1:500, ab109250, abcam; TRA-1-60, 1:500, ab109884, abcam; OCT4, 1:500, ab109884, abcam; SOX2, 1:500, ab109884, abcam; SSEA4, 1:500, ab109884, abcam; FOXA2, 1:500, ab10822, abcam). For indirect immunostaining, samples were washed x3 times with PBS and incubated for 2 h at room temperature with corresponding secondary antibody (α-mouse. 488, 1:500, ab150077, abcam; α-rabbit. 488,1:500, ab150077, abcam; α-goat. 594, 1:500, ab150077, abcam; α-rabbit. 647, 1:500, ab150075, abcam). Nuclear and actin counterstains were performed using Abcam iFluor conjugated phalloidin (ab176753, ab176759), and Hoechst 33342 at 0.1 µg/mL (Sigma, 14533) dilute in x1 PBS and incubated for 30 mins at room temperature. Following counterstain incubation, samples were washed an additional x3 with PBS containing 0.05% w/v % sodium azide. For YAP and RUNX2 translocation studies, microscopy was completed on a Zeiss LSM 800 Confocal microscope using 63x Objective Plan-Apochromat 63x/1.40NA Oil objective (with an in-plane lateral resolution of 0.413–0.124 µm per pixel) and pinhole diameter of 1.0 AU and azimuthal resolution of 0.4 µm. Nuclear images from Hoechst staining were used to create masks that define a nuclear volume. hADSC cytoplasmic masks were defined from flood-filled phalloidin stains, with the average fluorescent intensity of each volume calculated in MATLAB. Therein, the YAP/RUNX2 nuclear to cytoplasmic translocation ratio was determined as the ratio of the mean YAP/RUNX2 fluorescent saturation intensity of the nuclear volume divided by the fluorescent saturation intensity in the non-nuclear cell cytoplasmic volume. In **Fig. 3b,c, Fig. 4a,c**, YAP/RUNX2 measurements of a total of at least 12 single cells per condition were pooled across 3 independent experimental replicates from at least 4 printed-niche-replicates. Representative images were selected according to their proximity to the mean data as calculated across all replicates. For imaging hiPSCs, we used a Zeiss LSM 800 Confocal microscope using a Plan-Apo 20x/0.8NA. Using Hoechst nuclear marker, individual cell nuclei were segmented using the StarDist algorithm^60,61^, defining nuclear masks. Using the nuclei masks, the fluorescent intensity of each stain channel was calculated. Then the coordinates of each nucleus within the niches were calculated, allowing replicate data to be remapped to a single plot that showed the average position mapped immunostained expression of the markers with marker-size, height, and opacity scaled proportionally to the fluorescent intensity of the nuclei (see SI for additional details). All experiments containing expression mapped markers were completed from at least 3 independent experimental replicates, except for experiments analyzing the effects of different concentrations of pSMAD1 (**Fig. 6e-g, S20-21**) completed in 2 independent replicates.

### LipidTOX, CNA35, and Alizarin Red staining of hADSC differentiation

We assayed the differentiation of hADSCs toward adipogenic and osteogenic lineage in response to niche microproperties. All solutions listed below were syringe filtered through 0.22 µm membrane filters (Merck Millipore SLGP033RS). Alizarin Red (Sigma, A5533) staining was performed to examine the presence of mineralized deposits under osteogenic differentiation conditions. Samples were washed x2 with PBS before fixation at room temperature in 4% PFA dilute in x1 PBS buffer for 10 mins and then washed three times with PBS. Samples were then washed x3 in Milli-Q H2O before incubation with Alizarin Red stain for 5 minutes (9.6 mg/mL Alizarin Red at a pH of 4.2, adjusted with acetic acid). Following incubation, samples were washed x5 with Milli-Q H2O, followed by a further x3 washes with x1 PBS containing 0.05 w/v % sodium azide. We examined cell mineralization with Alizarin Red staining and widefield Color microscopy of materials following 14 d of culture. Cells were imaged using a Nikon Ni-E microscope with color DS-Fi2 camera and Plan Apo Lambda 10x/0.45NA dry objective. The localization of osteogenesis over specified RGD and Young’s modulus regions was quantified for each niche-replicate sample. The mean red saturation was divided by the mean total saturation that combines red, green, and blue color components for the corresponding region of interest. For Alizarin Red Stains in **Fig. 4b,d,h,k**, measurements were pooled across 3 independent experimental replicates from at least 4 printed-niche-replicates in each experiment for at least 12 total measurements per niche condition. Staining with LipidTOX Red Neutral Lipid Stain (Thermo H34476) was completed to quantify cell fat volume under adipogenic differentiation conditions. Samples were fixed at room temperature 4% PFA in x1 PBS buffer at pH 7.4 for 10 mins and then washed three times with PBS, followed by 12 min permeabilization at room temperature with 0.1 w/v % Triton X-100 in PBS. Samples were then incubated with LipidTOX Red Neutral Lipid Stain (diluted 1:800), Abcam iFluor conjugated phalloidin (1:200), and Hoechst 33342 at 0.1 µg/mL (Sigma, 14533) dilute in x1 PBS for 30 mins at room temperature. For LipidTOX data in **Fig. 4e-f**, measurements were pooled across 3 independent experimental replicates from at least 4 printed-niche-replicates in each experiment for a total of at least 24 fields-of-view per niche condition. In **Fig. 4k**, LipidTOX measurements were pooled across 3 independent experimental replicates from at least 4 printed-niche-replicates in each experiment for a total of 12 bone-fat niches. Following incubation, samples were washed an additional x3 with x1 PBS containing 0.05 w/v % sodium azide. Fluorescent microscopy of large fields of view (arrays of RGD and Young’s modulus) were tiled using a Nikon Ni-E microscope with a motorized stage, monochrome DS-Qi2 camera and Plan Apo Lambda 10x/0.45NA dry objective. Confocal microscopy was completed on a Zeiss LSM 800 Confocal microscope using x63 Plan-Apo 63x/1.40NA Oil objective (with an in-plane lateral resolution of 0.413 µm per pixel), pinhole diameter of 1.0 AU (50.34 µm), and azimuthal resolution of 0.4 µm. Segmentation was performed using custom MATLAB scripting that makes use of the open microscopy Bio-Formats tool. Masks were created to define cytoplasmic and fat volumes using phalloidin and LipidTOX stains. For production and purification of the fluorescent collagen 1A probe, the pET28a-EGFP-CNA35 plasmid was received as a gift from Maarten Merkx (Addgene plasmid # 61603; http://n2t.net/addgene:61603; RRID: Addgene_61603) and synthesized as reported previously^62^. In brief, protein yields of the CNA35 probe were synthesized using E.Coli bacteria before purification using ÄKTApurifier (Cytiva) and a 5 ml Ni-NTA Superflow Cartridge (Qiagen), dialysis with SnakeSkinTM Dialysis tubing with 10 kDa MWCO, and concentration with an Amicon 10 kDa MWCO centrifugal filter unit. For imaging, 0.5 µM of EGFP-CNA35 solution was added to the sample and incubated on a plate rocker for 15 mins, before washing twice with PBS. In **Fig. 4k**, Col1A measurements were pooled across 3 independent experimental replicates from at least 4 printed-niche-replicates in each experiment for a total of 12 bone-fat niches.

## Acknowledgments

The authors gratefully acknowledge the Australian Government’s financial support for providing PLHN an Australian Postgraduate Award Scholarship, the Australian Department of Education and Training for awarding PLHN with an Endeavour Fellowship. A massive thank you to the discussions and support of Dr. Martin Stewart, Dr. Ashnil Kumar, Dr. Anai Gonzalez-Cordero and Hélène Lebhar from the UNSW Recombinant Products Facility for assistance in the production and purification of EGFP-CNA35. From Dr. Courtney Wright, aka the mensch, the facilities and the scientific and technical aid of Microscopy Australia at the Australian Centre for Microscopy & Microanalysis at the University of Sydney, including Dr. Ying Ying Su and Dr. Neftali Flores-Rodriguez. PN thanks Karl Gifford for his critical assistance in discussing and building custom 3D printers.

## Author contributions

PN conceived the work, designed the modified MCFL 3D-printer device and method, designed and executed all experimentation, analyzed all data, and wrote the manuscript. QY and PO assisted manuscript preparation, and together with JS, helped the development of methods relating to the hiPSCs. TA assisted with printing structures for hADSCs. DK prepared and provided CNA fluorescent probes and methods relating to their use. MB provided scientific oversight, experimental analysis and assisted with manuscript preparation. PT funded experiments relating to hiPSCs, assisted experimental design and methods on the culture of hiPSCs, provided scientific oversight, and assisted with manuscript preparation. J-WS funded and provided scientific oversight developing methods for specifying niche mechanics and related force spectroscopic measurement. HZ funded the project, provided scientific oversight, and assisted with manuscript preparation.

## Data and materials availability

Standalone software or software tools were developed for this work and provided as supplemental materials, with up-to-date software available by request to lead authors. All scripts used for data analysis are available upon request to the corresponding authors. Datasets supporting the conclusions are shown within the article and its additional files. Data and related scripts with sample data for analysis are available upon reasonable request to the corresponding authors.

## References

1. Collinet, C. & Lecuit, T. Programmed and self-organized flow of information during morphogenesis. Nat. Rev. Mol. Cell Biol. 1–21 (2021).

2. Hannezo, E. & Heisenberg, C.-P. Mechanochemical feedback loops in development and disease. Cell 178, 12–25 (2019).

3. Warmflash, A., Sorre, B., Etoc, F., Siggia, E. D. & Brivanlou, A. H. A method to recapitulate early embryonic spatial patterning in human embryonic stem cells. Nat. Methods 11, 847 (2014).

4. Zheng, Y. et al. Controlled modelling of human epiblast and amnion development using stem cells. Nature 573, 421–425 (2019).

5. Nikolaev, M. et al. Homeostatic mini-intestines through scaffold-guided organoid morphogenesis. Nature (2020) doi:10.1038/s41586-020-2724-8.

6. Birey, F. et al. Assembly of functionally integrated human forebrain spheroids. Nature 545, 54–59 (2017).

7. Engler, A. J., Sen, S., Sweeney, H. L. & Discher, D. E. Matrix elasticity directs stem cell lineage specification. Cell 126, 677–689 (2006).

8. Cameron, A. R., Frith, J. E. & Cooper-White, J. J. The influence of substrate creep on mesenchymal stem cell behaviour and phenotype. Biomaterials 32, 5979–5993 (2011).

9. Dalby, M. J., Gadegaard, N. & Oreffo, R. O. Harnessing nanotopography and integrin– matrix interactions to influence stem cell fate. Nat. Mater. 13, 558–569 (2014).

10. Khetan, S. et al. Degradation-mediated cellular traction directs stem cell fate in covalently crosslinked three-dimensional hydrogels. Nat. Mater. 12, 458–465 (2013).

11. Hosaka, S., Ozawa, H. & Tanzawa, H. Controlled release of drugs from hydrogel matrices. J. Appl. Polym. Sci. 23, 2089–2098 (1979).

12. Edelman, E. R., Mathiowitz, E., Langer, R. & Klagsbrun, M. Controlled and modulated release of basic fibroblast growth factor. Biomaterials 12, 619–626 (1991).

13. Burdick, J. A. & Anseth, K. S. Photoencapsulation of osteoblasts in injectable RGD-modified PEG hydrogels for bone tissue engineering. Biomaterials 23, 4315–4323 (2002).

14. Cosgrove, B. D. et al. N-cadherin adhesive interactions modulate matrix mechanosensing and fate commitment of mesenchymal stem cells. Nat. Mater. 15, 1297– 1306 (2016).

15. Miller, J. S. et al. Rapid casting of patterned vascular networks for perfusable engineered three-dimensional tissues. Nat. Mater. 11, 768–774 (2012).

16. Xu, T. et al. Complex heterogeneous tissue constructs containing multiple cell types prepared by inkjet printing technology. Biomaterials 34, 130–139 (2013).

17. Hockaday, L. et al. Rapid 3D printing of anatomically accurate and mechanically heterogeneous aortic valve hydrogel scaffolds. Biofabrication 4, 035005 (2012).

18. Kolesky, D. B., Homan, K. A., Skylar-Scott, M. A. & Lewis, J. A. Three-dimensional bioprinting of thick vascularized tissues. Proc. Natl. Acad. Sci. 113, 3179–3184 (2016).

19. Kang, H.-W. et al. A 3D bioprinting system to produce human-scale tissue constructs with structural integrity. Nat. Biotechnol. 34, 312–319 (2016).

20. Ober, T. J., Foresti, D. & Lewis, J. A. Active mixing of complex fluids at the microscale. Proc. Natl. Acad. Sci. 112, 12293–12298 (2015).

21. Skylar-Scott, M. A., Mueller, J., Visser, C. W. & Lewis, J. A. Voxelated soft matter via multimaterial multinozzle 3D printing. Nature 575, 330–335 (2019).

22. Brassard, J. A., Nikolaev, M., Hübscher, T., Hofer, M. & Lutolf, M. P. Recapitulating macro-scale tissue self-organization through organoid bioprinting. Nat. Mater. (2020) doi:10.1038/s41563-020-00803-5.

23. Lee, A. et al. 3D bioprinting of collagen to rebuild components of the human heart. Science 365, 482–487 (2019).

24. Kolesky, D. B. et al. 3D bioprinting of vascularized, heterogeneous cell-laden tissue constructs. Adv. Mater. 26, 3124–3130 (2014).

25. Richter, B. et al. Guiding cell attachment in 3D microscaffolds selectively functionalized with two distinct adhesion proteins. Adv. Mater. 29, 1604342 (2017).

26. Ma, X. et al. Deterministically patterned biomimetic human iPSC-derived hepatic model via rapid 3D bioprinting. Proc. Natl. Acad. Sci. 113, 2206–2211 (2016).

27. Lawlor, K. T. et al. Cellular extrusion bioprinting improves kidney organoid reproducibility and conformation. Nat. Mater. 1–12 (2020).

28. Müller, P. et al. STED-inspired laser lithography based on photoswitchable spirothiopyran moieties. Chem. Mater. 31, 1966–1972 (2019).

29. Chen, C. S., Mrksich, M., Huang, S., Whitesides, G. M. & Ingber, D. E. Geometric control of cell life and death. Science 276, 1425–1428 (1997).

30. Mosiewicz, K. A. et al. In situ cell manipulation through enzymatic hydrogel photopatterning. Nat. Mater. 12, 1072–1078 (2013).

31. Luo, Y. & Shoichet, M. S. A photolabile hydrogel for guided three-dimensional cell growth and migration. Nat. Mater. 3, 249–253 (2004).

32. Shadish, J. A., Benuska, G. M. & DeForest, C. A. Bioactive site-specifically modified proteins for 4D patterning of gel biomaterials. Nat. Mater. 18, 1005–1014 (2019).

33. Yin, H., Ding, Y., Zhai, Y., Tan, W. & Yin, X. Orthogonal programming of heterogeneous micro-mechano-environments and geometries in three-dimensional bio-stereolithography. Nat. Commun. 9, 1–7 (2018).

34. Mayer, F. et al. Multimaterial 3D laser microprinting using an integrated microfluidic system. Sci. Adv. 5, eaau9160 (2019).

35. Dendukuri, D., Pregibon, D. C., Collins, J., Hatton, T. A. & Doyle, P. S. Continuous-flow lithography for high-throughput microparticle synthesis. Nat. Mater. 5, 365–369 (2006).

36. Fu, J., Warmflash, A. & Lutolf, M. P. Stem-cell-based embryo models for fundamental research and translation. Nat. Mater. 1–13 (2020).

37. Rossant, J. & Tam, P. P. Emerging asymmetry and embryonic patterning in early mouse development. Dev. Cell 7, 155–164 (2004).

38. Rosado-Olivieri, E. A. & Brivanlou, A. H. Synthetic by design: exploiting tissue self-organization to explore early human embryology. Dev. Biol. 474, 16–21 (2021).

39. Cornwall-Scoones, J. & Zernicka-Goetz, M. Unifying synthetic embryology. Dev. Biol. 474, 1 (2021).

40. Hofer, M. & Lutolf, M. P. Engineering organoids. Nat. Rev. Mater. 6, 402–420 (2021).

41. Valet, M., Siggia, E. D. & Brivanlou, A. H. Mechanical regulation of early vertebrate embryogenesis. Nat. Rev. Mol. Cell Biol. 1–16 (2021).

42. Sheng, G., Martinez Arias, A. & Sutherland, A. The primitive streak and cellular principles of building an amniote body through gastrulation. Science 374, abg1727 (2021).

43. Fairbanks, B. D., Schwartz, M. P., Bowman, C. N. & Anseth, K. S. Photoinitiated polymerization of PEG-diacrylate with lithium phenyl-2, 4, 6-trimethylbenzoylphosphinate: polymerization rate and cytocompatibility. Biomaterials 30, 6702–6707 (2009).

44. Guimarães, C. F., Gasperini, L., Marques, A. P. & Reis, R. L. The stiffness of living tissues and its implications for tissue engineering. Nat. Rev. Mater. 1–20 (2020).

45. Dupont, S. et al. Role of YAP/TAZ in mechanotransduction. Nature 474, 179–183 (2011).

46. Caliari, S. R., Vega, S. L., Kwon, M., Soulas, E. M. & Burdick, J. A. Dimensionality and spreading influence MSC YAP/TAZ signaling in hydrogel environments. Biomaterials 103, 314–323 (2016).

47. Meng, Z., Moroishi, T. & Guan, K.-L. Mechanisms of Hippo pathway regulation. Genes Dev. 30, 1–17 (2016).

48. Chaudhuri, O. et al. Hydrogels with tunable stress relaxation regulate stem cell fate and activity. Nat. Mater. 15, 326–334 (2016).

49. Komori, T. Regulation of bone development and extracellular matrix protein genes by RUNX2. Cell Tissue Res. 339, 189 (2010).

50. Bianco, P., Robey, P. G. & Simmons, P. J. Mesenchymal stem cells: revisiting history, concepts, and assays. Cell Stem Cell 2, 313–319 (2008).

51. Chan, C. K. et al. Identification of the human skeletal stem cell. Cell 175, 43–56 (2018).

52. Mongera, A. et al. A fluid-to-solid jamming transition underlies vertebrate body axis elongation. Nature 561, 401–405 (2018).

53. Nakaya, Y., Sukowati, E. W., Wu, Y. & Sheng, G. RhoA and microtubule dynamics control cell–basement membrane interaction in EMT during gastrulation. Nat. Cell Biol. 10, 765–775 (2008).

54. Kyprianou, C. et al. Basement membrane remodelling regulates mouse embryogenesis. Nature 1–6 (2020).

55. Schmidt, U., Weigert, M., Broaddus, C. & Myers, G. Cell detection with star-convex polygons. in 265–273 (Springer, 2018).

56. Weigert, M., Schmidt, U., Haase, R., Sugawara, K. & Myers, G. Star-convex polyhedra for 3d object detection and segmentation in microscopy. in 3666–3673 (2020).

57. Simunovic, M. et al. A 3D model of a human epiblast reveals BMP4-driven symmetry breaking. Nat. Cell Biol. 21, 900–910 (2019).

58. Chhabra, S., Liu, L., Goh, R., Kong, X. & Warmflash, A. Dissecting the dynamics of signaling events in the BMP, WNT, and NODAL cascade during self-organized fate patterning in human gastruloids. PLoS Biol. 17, e3000498 (2019).

59. Liu, X. et al. Modelling human blastocysts by reprogramming fibroblasts into iBlastoids. Nature 591, 627–632 (2021).

60. Muncie, J. M., Ayad, N. M., Lakins, J. N. & Weaver, V. M. Mechanics regulate human embryonic stem cell self-organization to specify mesoderm. Dev.-CELL--20-00131 (2020).

61. Tewary, M. et al. High-throughput micropatterning platform reveals Nodal-dependent bisection of peri-gastrulation–associated versus preneurulation-associated fate patterning. PLoS Biol. 17, e3000081 (2019).

62. Grau, Y., Carteret, C. & Simpson, P. Mutations and chromosomal rearrangements affecting the expression of snail, a gene involved in embryonic patterning in Drosophila melanogaster. Genetics 108, 347–360 (1984).

63. Xu, P.-F., Houssin, N., Ferri-Lagneau, K. F., Thisse, B. & Thisse, C. Construction of a vertebrate embryo from two opposing morphogen gradients. Science 344, 87–89 (2014).

64. Manfrin, A. et al. Engineered signaling centers for the spatially controlled patterning of human pluripotent stem cells. Nat. Methods 16, 640 (2019).

65. Rivron, N. C. et al. Blastocyst-like structures generated solely from stem cells. Nature 557, 106–111 (2018).

66. Kagawa, H. et al. Human blastoids model blastocyst development and implantation. Nature 1–9 (2021).

67. Izquierdo, E., Quinkler, T. & De Renzis, S. Guided morphogenesis through optogenetic activation of Rho signalling during early Drosophila embryogenesis. Nat. Commun. 9, 1–13 (2018).

68. Grigoryan, B. et al. Multivascular networks and functional intravascular topologies within biocompatible hydrogels. Science 364, 458–464 (2019).

69. Wylie, R. G. et al. Spatially controlled simultaneous patterning of multiple growth factors in three-dimensional hydrogels. Nat. Mater. 10, 799–806 (2011).

