## Supplemental Materials for "Material-mediated histogenesis using mechano-chemically microstructured cell niches"

**Overview of printer.** Details of the custom-built mechano-chemical flow lithographic (MCFL) printer are provided below including the printing method in-line explanatory figures (**Fig. S 1**). This includes fabricating one-time-use glass components, calibrating the printer, loading photoresists, printing, software use, and preparing samples for cell seeding. All components of the printer are detailed in **Table S 1**.

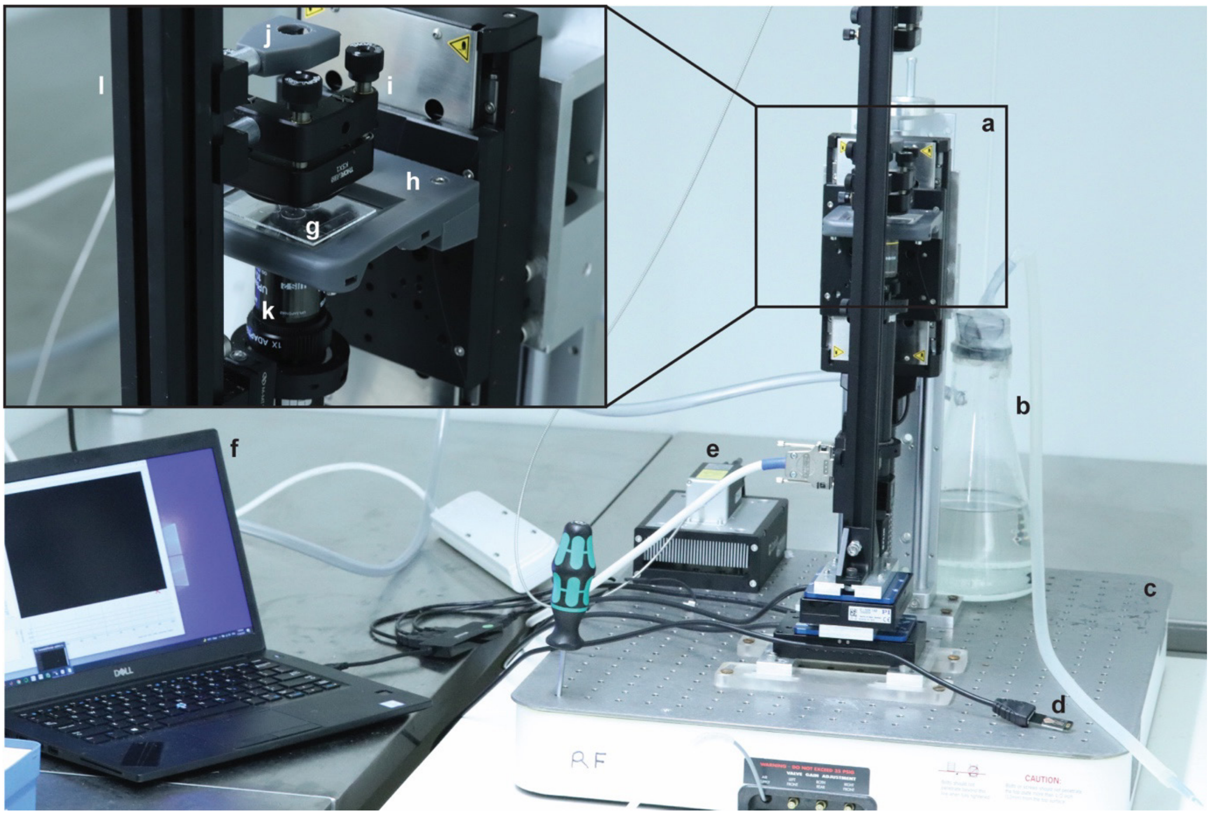

**Fig. S 1 | Overview and labeled setup of the MCFL printer. a,** MCFL printer stage. **b,** vacuum system. **c,** optical breadboard table. **d,** LED touch light. **e,** diode laser and heatsink. **f,** computer with software. The inset at the top left shows a detailed view of the print chamber. **g,** glass sample chamber. **h,** custom-printed stage. **i,** mounted aspheric lens. **l,** optical rail. **j,** custom-printed LED housing. **k,** objective lens.

Niches were fabricated on glass 10 mm diameter size #1 coverslips by the injection or 'flowing' of photoresists of variable biochemical and polymeric composition through a print chamber during printing (**Fig. 1**, **Fig. S1h,** see **Table S 2** for specific photoresist compositions). **Fig. S 2** shows an isolated view of the printer stage platform, alongside the PDMS coated glass slide loaded with four coverslips.

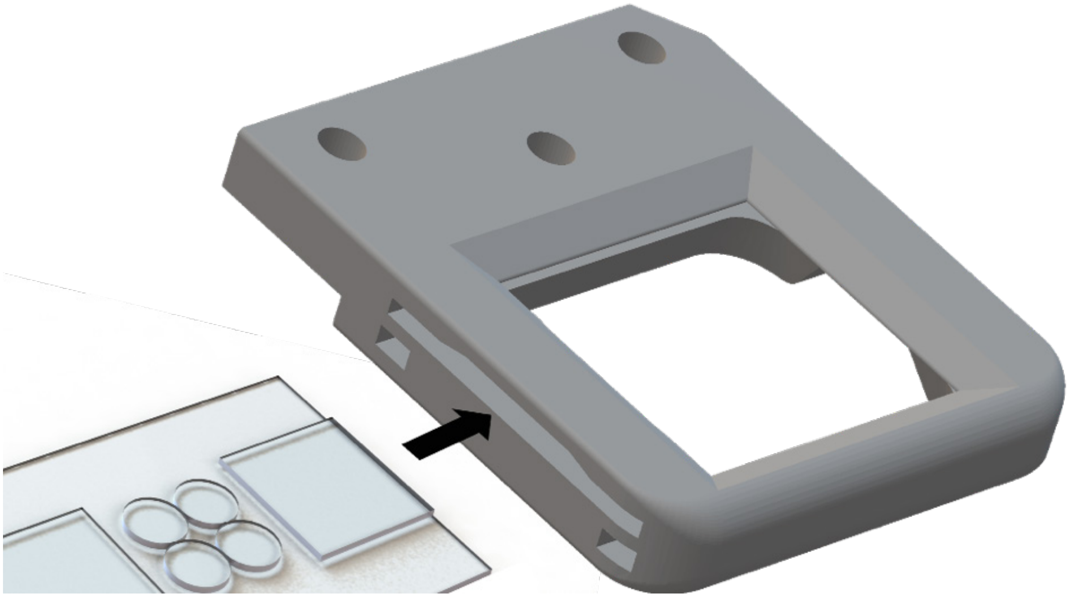

**Fig. S 2 | Schematic of custom-printed stage housing for printing chamber.** The constructed printing chamber base is inserted into the side slot and secured through clamping bolts under each corner. A photoresist is injected onto the 10 mm acrylated coverslips, and a PDMS-coated top slide is positioned centrally over the opening.

**Fabrication of print chamber and sample coverslips.** A photoresist is injected onto 10 mm diameters coverslips sandwiched by two hydrophobic PDMS coated glass slides during printing. This forms the print chamber (**Fig. S1-a,b**), is then clamped into the printer stage. PDMS coated 50x75x1 mm glass slides (Sigma CLS294775X50) are prepared using PDMS (Corning Sylgard 184) ten-parts base to one-part curing agent. The solution is mixed with a metal spatula and briefly centrifuged to remove bubbles. Glass slides are spin-coated with PDMS on a Laurell EDC 650 series spin coater. PDMS is poured onto the top of the samples at approximately an Australian 50 cent piece volume. The spin coater is then ramped to 1600 rpm for 10 s and stopped. The glass slides are then placed onto a hot plate at 200 ˚C for 1 min before being transferred to a 37 ˚C oven overnight.

Printing was completed on an acrylate-functionalized coverslip base that forms a stable substrate for transport and culture of the printed materials, including the prevention of sample delamination, sample-folding, or strain deformation due to hydrogel swelling, as well, the covalent attachment assists sample handling for downstream cell culture. A silanization solution was prepared to functionalize coverslips. A glass dish was washed with methanol (Sigma 322415) by bath sonicating for 5 minutes in a chemical safety fume hood. The dish was then dried then rinsed a further x2 with methanol. 100 mL of methanol, 5 mL of glacial acetic acid (Sigma A6283), and 3 mL 3-(trimethoxy silyl) propyl acrylate (TCI A1597) was added to the dish along. Circular glass coverslips were washed x3 times in methanol before being placed in the silanization solution. The dish was covered to prevent evaporation or contamination from ambient H2O. Coverslips are left for one hour for the silanization reaction to proceed. Coverslips were then rinsed x3 in ethanol, wiped clean, and dried with $N_2$. All glass and PDMS components are thoroughly washed with warm tap water, Triton X-100, acetone, isopropanol, and purified water once again prior to printing. The glass is then air-dried using high-pressure filtered compressed air to limit dust particles and lint on samples.

The print chamber is assembled by suctioning the acrylated coverslips onto the PDMS-coated glass slide. A single print may consist of four 10 mm coverslips positioned in the middle of the slide **(Fig. S 2)**. Suction force holds sample coverslips to the PDMS coating. Plain square glass coverslips (24 mm x 24 mm) are positioned adjacent to each short edge of the PDMS slide to ensure the correct elevation of the chamber's top **(Fig. S 2)**. The top chamber uses an opposing PDMS glass slide and effectively sandwiches the acrylated substrate coverslips within a hydrophobic glass chamber when positioned on top. The printer setup is housed within a dark 4°C cold room pictured throughout - to limit photoactivation of the resist, as well as damage, misfolding, and gelling of sensitive biochemical reagents. All printer components and reagents are stored in the cold room to acclimatize before printing, preventing condensation. The base PDMS slide is inserted into the printing stage and secured into position through bolts at the bottom of each corner **(Fig. S 2)**. The chamber's top is then positioned to calibrate coverslips and chamber height. Machine bolts then attach the printer stage to the rest of the printer.

**Calibration of printer laser with sample coverslips and stage.** A camera assembly is mounted below the printing stage to monitor laser alignment and focus. The printer's laser is first calibrated to the camera assembly (**Table S 1** for component details) using the custom printer software (**Fig. S 3**). The distance between the laser and camera assembly is altered for parfocality. The camera is focused on the stage by changing the stage height. The laser is then focused through a position that passes the acrylated coverslip by navigation using the GUI stage map within the printer control panel (**Fig. S 3c**). Focusing the laser at this position ensures an accurate focal length of the light path that includes the refractive differences of all materials and prevents aberrations caused by coverslip edges. During this step, focal asymmetry and astigmatisms can be reduced by carefully calibrating machine thumbscrews on mechano-optical components housing the laser fiber unit, collimator, beam expander, and focal objective (for parts numbers see **Table S 1**).

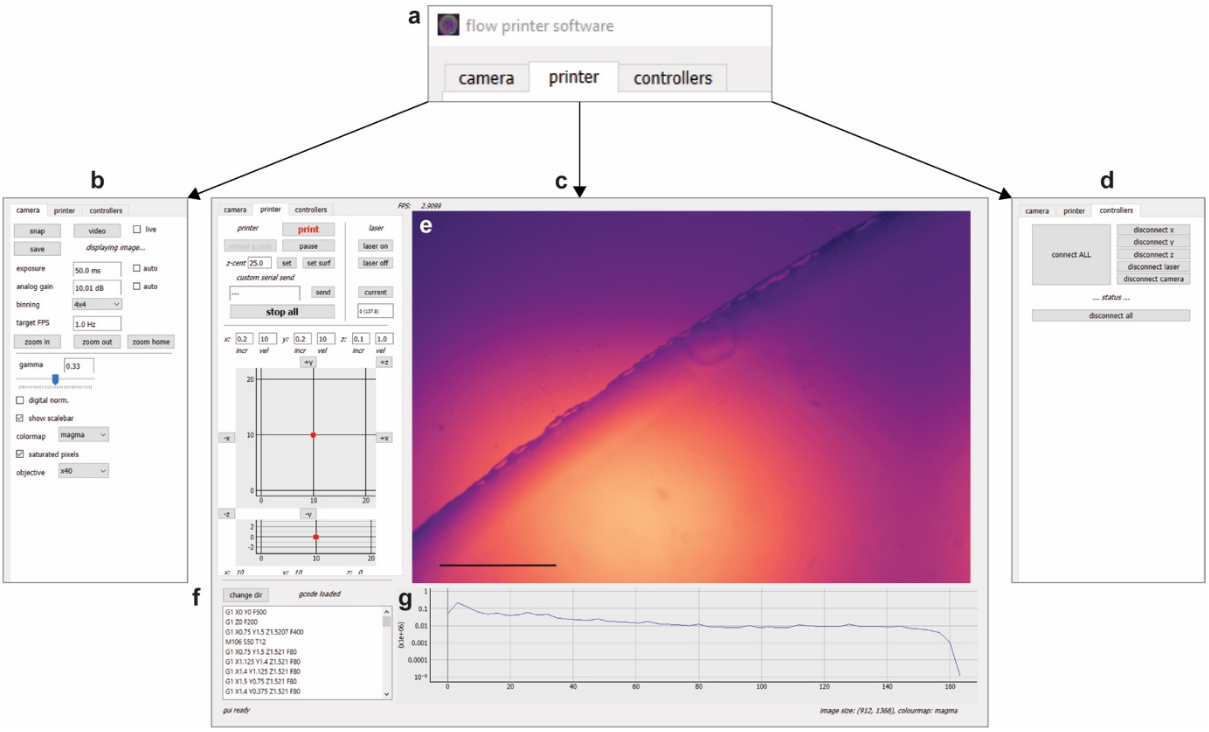

**Fig. S 3 | Labelled layout of software for printer operation.** Three panels are available to control printer parameters; camera, printer, controllers. **a,** Software interface panels. **b,** camera control. **c,** printer control. **d,** printer connection diagnostics with controllers. **e,g** live camera image output (pictured at the calibrated edge of coverslip sample). **f,** gcode input/output, and control. Standalone printer software as written in python with PyQt5 is available for download from supplemental information.

The stage height is adjusted to focus on the print sample during laser calibration, ensuring calibration between the laser, camera, and stage/samples. Camera parameters such as exposure, gain, binning, frames per second (FPS), and gamma may be adjusted in the GUI to visualize the live image better. A further feature of the camera view includes toggles for colormap, scalebar, digital normalization, saturated pixels, and objective may also be toggled for viewing experience **(Fig. S 3b)**.

**Injection of photoresist into glass print chamber.** Following calibration of the printer, a photoresist is injected within the print chamber onto the acrylated coverslips. Accurate assembly of the print chamber results in a 50 µm void between the PDMS top slide and sample for photoresist injection.

**gcode upload and print initiation.** The software interprets and translates gcode commands, sending these to the PI controllers - (V-528.1AA /V-528.1AB and M-406 including corresponding controllers C-413 and C863 from Physik Instrumente (PI) GmbH & Co. KG). The gcode is loaded by navigating to the relevant directory and then visually verified through the log on the bottom left of the GUI **(Fig. S 3f)**. An inspection of the printing area is performed to ensure no objects are obstructing printer movement. Printing is initiated and confirmed for correct laser movement. If errors are present, printing may be paused or terminated in the GUI. When performing a print with multiple photoresists, the vacuum pump is used to suction any excess photoresist from the previous print. The coverslips are gently washed with PBS supplemented with penicillin and streptomycin. If a printer connection is lost during the process, this may be troubleshot using the controller tab. Here, the connection to the stage, laser, and camera may be disconnected or reconnected. Otherwise, the printer may be switched on and off to restore the connection.

**Removal of samples from the printer.** The top slide is carefully removed after printer termination to ensure that the printed substrate is not damaged or attached to the chamber. The stage is then unfastened from the printer and removed. The printer chamber is unclamped and taken out of the stage, revealing the samples. For each coverslip, a well of a 48-well plate is prepared with 0.5 mL of PBS supplemented with penicillin and streptomycin. Coverslips are removed from the base PDMS-coated glass using a scalpel blade to gently disturb the suction forces between the sample base and the PDMS. Care is taken not to damage the printed substrate or glass coverslip. The coverslips are then moved to a well, ensuring that the printed substrate faces the right side upwards. Well plates containing samples should then be parafilmed and left to wash on a plate rocker at 4 ˚C for 48h before use. Samples have been stored for up to 1 week without observing any changes to their function.

**Sterilization and preparation of printed substrates.** The printed niches are washed two times with sterile PBS prior to cell seeding. While submerged, samples are UV-sterilized in a biosafety cabinet for 12 minutes, with the plate lid removed. For circular glass control samples, coverslip samples are incubated with hESC qualified Matrigel® (Corning) for 1 hour before cell seeding.

**3D structures in Fig. 1.** The printed structures presented in **Fig 1h,i** were achieved as per Grigoryan et al.[1], adding 3 mM of the photoabsorber tartrazine, as to limited the penetration depth of the laser . A full optimization of the variable state-space for MCFL of 3D structures using the tartrazine methods was unexplored in the present study. As per discussions points, this approach may permit further exploration of the method using complex 3D structures.

**Table S 1. Printer parts and components.** Labels column references item locations are annotated over **Fig. S 1**.

| Category | Description | Label |
|---|---|---|
| Laser Mount | RC4 - Dovetail Rail Carrier, 0.60" x 1.00" (15.2 mm x 25.4 mm), #8 (M4) Counterbore | |
| | K5X1 - 5-Axis Locking Kinematic Mount for Ø1" Optics | |
| Laser 2 | Cobolt 06-01 Series 405 nm, fiber pigtailed, FC/APC | S1e |
| | HS-03 Heat Sink | |
| | 405 nm FC/APC Collimation Package, NA = 0.26, f = 33.9 | |
| | K5X1 - 5-Axis Locking Kinematic Mount for Ø1" Optics | |
| | RC4 - Dovetail Rail Carrier, 0.60" x 1.00" (15.2 mm x 25.4 mm), #8 (M4) Counterbore | |
| Aspheric Lens | RC4 - Dovetail Rail Carrier, 0.60" x 1.00" (15.2 mm x 25.4 mm), #8 (M4) Counterbore | S1a, S1i |
| | KM100T - SM1-Threaded Kinematic Mount for Thin Ø1" Optics | |
| | S1TM12 - SM1 to M12 x 0.5 Lens Cell Adapter | |
| | A240TM - f = 8.0 mm, NA = 0.50, Mounted Rochester Aspheric Lens, Uncoated | |
| Optic Rail | XE25L900/M - 25 mm Square Construction Rail, 900 mm Long, M6 Taps | S1l |
| | RLA300/M - Dovetail Optical Rail, 300 mm, Metric | |
| | RLA300/M - Dovetail Optical Rail, 300 mm, Metric | |
| | XE25L225/M - 25 mm Square Construction Rail, 225 mm Long, M6 Taps | |

| | | |
|---|---|---|
| Sample/Slide Holder | KM100C - Kinematic Mount for up to 1.3" (33 mm) Tall Rectangular Optics, Right Handed | S1h, S2 |
| | Manual XYZ trimming stage for placing transparent stage | |
| | Z stage with sample holder Fig S1,2 | |
| Stage 3 10nm | M-406.2DG Precision Linear Stage | S1 |
| | V-528.1AA High-Dynamics PIMag® Linear Stage | |
| | V-528.1AB High-Dynamics PIMag® Linear Stage | |
| | C-413.2G PIMag® Motion Controller | |
| | C-863 Mercury Servo Controller | |
| Camera Assembly | RC4 - Dovetail Rail Carrier, 0.60" x 1.00" (15.2 mm x 25.4 mm), #8 (M4) Counterbore | S1k |
| | SM30TC - Clamp for SM30 Lens Tubes | |
| | x20 Olympus Objective - UPLSAPO10X2 | |
| | RMSA7 - Adapter with External M26 x 0.706 Threads and Internal RMS Threads | |
| | CMV10 - C-Mount Adjustable Extension Tube | |
| | MT-4 Accessory Tube Lens | |
| | Mitutoyo to C-mount Camera 152.5mm Extension Tube | |
| | NF405-13 - Ø25 mm Notch Filter, CWL = 405 nm, FWHM = 13 nm | |
| | C-Mount Filter/Polarizer Holder, M6 | |
| | 25/25.4mm Optic Cell | |
| | BFS-U3-200S6M-C: 20 MP, 18 FPS, SONY IMX183, MONO | |
| | ACC-01-2304: USB3, 1 M, TYPE-A TO MICRO-B LOCKING CABLE | |

| | |
|---|---|
| Breadboard | Newport Optical Breadboard Table BT-2024 24"x20"x3" |
| Custom-printed Components (Available via a request to lead authors) | Printing Sample Stage |
| | M406 Base Plate |
| | M406 Mount |
| | Rail Spacer |
| | Stage Clip |
| | Stage Mount |
| | V528 Base Plate |
| | V528 Mid Plate |
| | V528 Rail Mount |

**Changing focal distance for minimal linewidth.** The below image shows fluorescent microscopy of MCFL filaments of varying linewidth (printed with a 20 v v$^{-1}$ % PEGDA, 0.05 mg mL$^{-1}$ LAP, TRITC 0.56 mM solution, scan vel. 100 mm min$^{-1}$, duty-cycle/PWM 100). Filaments of varying linewidth were achieved through changes to the focus. A minimal linewidth of approximately 7 µm was achieved.

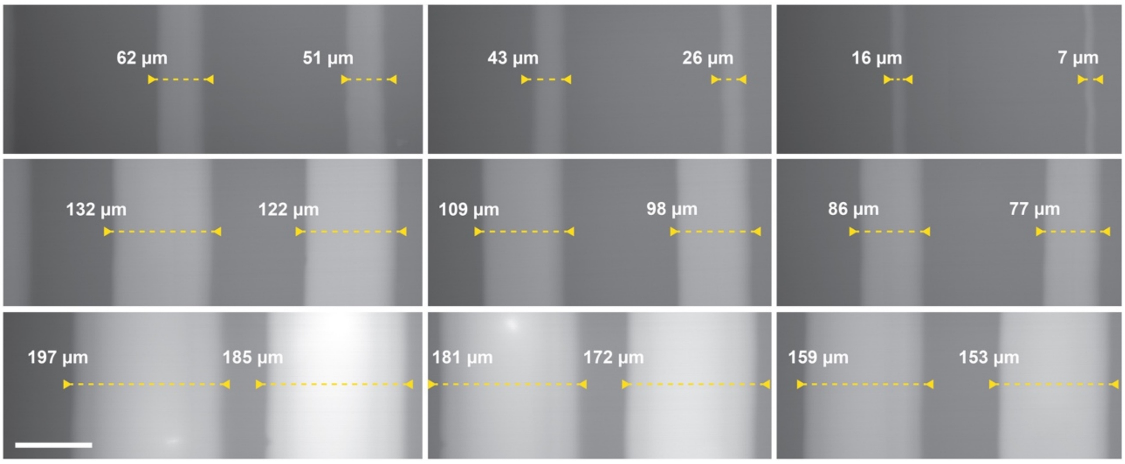

**Fig. S 4 | Changing focus for altering linewidth using MCFL 3DP.**

**Attachment assay with bulk hydrogels for various thiol-ene conjugated ligands and cell types.** The broader potential of the thiol-ene chemistry approach was initially tested using different ligands and cell types, including human bone marrow-derived mesenchymal stromal cells (hBDMSCs) and HUVECs (human vascular endothelial cells). The potential of this method was confirmed with bulk hydrogel disks cured with a UV flood lamp using the photoresist composition used throughout.

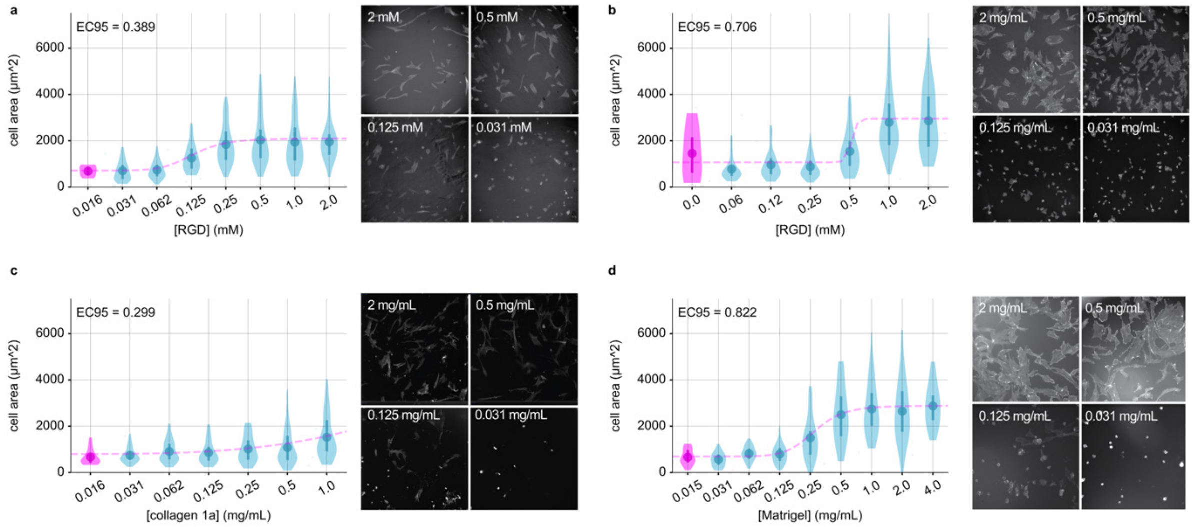

**Fig. S 5 | The thiol-ene bioconjugation approach was first broadly demonstrated for attachment ligands in bulk hydrogels.** Culturing both hBDMSCs and HUVECs for **a,** hBDMSCs with RGD attachment ligand **b,** HUVECs with RGD attachment ligand. **c,** hBDMSCs with rat tail collagen 1A attachment ligand**. d,** HUVECs with Matrigel attachment ligand. Representative images are shown for phalloidin staining.

**Notes on photopolymerization.** We demonstrated that increases to laser scan velocity decreased both Young's modulus and the linewidth of prints (**Fig. 2a**), as consistent with a lowered rate of polymerization due to the decreased light absorption and consequentially lower photoinitiator free radical dissociation (see notes below, **Fig. S 6, Fig. S 7**)[2]. Decreased laser scan velocity and hence increased light absorption are consistent with the effects of increasing the laser power during photopolymerization, where high power yielded increased Young's modulus and linewidth (**Fig. 2b**). As a function of the focus, changes to the linewidth were approximated with a linear function that correlated with the diameter of the conic angle of laser transmittance (**Fig. 2c**). The relationship between focus and Young's modulus is nonlinear, a behavior that may be related to photodamage of the niche or interference effects from imperfect optics. Increases in the photoinitiator and monomer concentration produced materials with higher Young's modulus and larger linewidths. This is consistent with accepted photopolymerization models that predict more polymer crosslinking as well as a larger volume exceeding the critical threshold to polymerization for photoresists with higher monomer and photoinitiator concentration (see notes below, **Fig. S 6**)[2].

Under irradiation, type I photoinitiators such as the LAP (Lithium phenyl-2,4,6-trimethylbenzoylphosphinate) photoinitiators dissociate into two or more radical species. For type I free radical photopolymerizing systems, a critical exposure threshold exists where a liquid photopolymer is solidified. This occurs when a photoresist absorbs sufficient light energy, such that free radicals generated from light-induced dissociation of a photoinitiator can overcome free radical inhibition (photoinitiator quenching) to crosslink a monomer and convert it to a solid. Increases to either the incident laser power (See **Fig. S 6b–f**) or laser residence time (**Fig. S 6d**) will result in a larger volume of polymerization. i.e., Other factors that alter the critical exposure threshold include changes to the concentration of photoinitiator, monomer, changes to the reaction atmosphere (changes to ambient [$O_2$] and absorbed oxygen radials), radical quenching agents such as glucose oxidase, photobleachers, photoabsorbers, and dyes. As can be observed in **Fig. S 6b,** changes to the critical exposure threshold result from altering system linewidth.

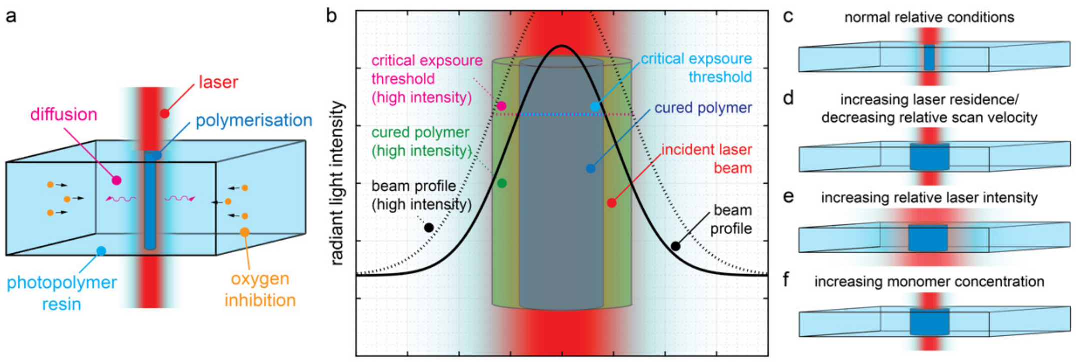

**Fig. S 6 | Shows a model photoresist during irradiance. a,** Key interactions are outlined, including laser irradiance and diffusive/oxygen inhibitory species. **b,** The critical exposure threshold and the effect on polymerization and linewidth are shown for varying laser irradiance. A high irradiance results in the polymerization of a larger volume of photoresist. **c,** The effect of varying irradiance is compared, showing changes to the polymerization volume.

We examined the effect of changing monomer concentration, photoinitiator concentration, scan velocity, incident light power, and light focus. The standard model of free radical photopolymerization expresses the rate of photopolymerization as[2]:

$$R_p = k_p[M]\left(\frac{\Phi\alpha[PI]I_0 10^3 e^{-\alpha[PI]D}}{k_t}\right)^{1/2} \quad (1)$$

This expression considers the rate of photopolymerization ($R_p$) as a function of monomer concentration ($[M]$), photoinitiator concentration ($[PI]$), and the absorbed light intensity ($I_0$) (where light power, light focus, and scan velocity all alter the absorbed light intensity). The remaining variables in Equation 1 above are ($D$) the depth of penetration, ($\alpha$) the absorption coefficient or molar absorptivity, ($\Phi$) the quantum yield for initiation or number of propagating chains initiated per light photon absorbed, ($k_p$) rate constant for polymer chain propagation and ($k_t$) termination rate constant. This model reveals a high sensitivity to changes in the photoinitiator concentration, as per the exponential term $e^{-\alpha[PI]D}$. Accordingly, uniformity in a photoresist is best achieved using as low of a concentration of a photoinitiator as possible, such that the exponent approximates 0. Thus, thin reaction systems ($D$ ~ 0) with low photoinitiator concentrations ($[PI]$ ~ 0) are reduced such that the exponent $\alpha[PI]D \sim 0$, and the expression reduces such that:

$$R_p \propto [M];\ R_p \propto {I_0}^{1/2};\ and\ R_p \propto [PI]^{1/2} \quad (2)$$

As the photopolymerization rate relates to the critical exposure threshold, so does it relate to the linewidth since polymerization occurs in this region. Changes to the scan velocity are inversely proportional to the absorbed light intensity (moving twice as fast would half the light exposure); this predicts that the linewidth and scan velocity should exhibit a ½-order relationship. However, fitting the data presented in **Fig. 2** with this model does not approximate the observed behavior (see **Fig. S 6a** below). Alternatively, fitting the data to an unknown order reveals the 0.28$^{th}$ order. This shows complex interactions that the model does not account for, suggesting that the criterion of low *[PI]* and *D* is not met for the MCFL device herein. Similar to changes in scan velocity, we fit the observed linewidth as a function of the photoinitiator and light power/duty-cycle. Fitting the data presented in **Fig. 2** with the above model better approximates the observed behavior (see **Fig. S 6b-c** below). Otherwise, fitting data to an unknown order reveals the photoinitiator concentration approximates a 0.61$^{th}$ order relationship with linewidth, and the laser duty-cycle approximates a 0.40$^{th}$ order relationship with linewidth. Considering the effects of diffusing radical species as well as changes to the concentration of inhibiting species such as that from ambient oxygen could help explain these observations. This analysis demonstrates that photoresist systems are highly sensitive, nonlinear, and difficult to predict. Accordingly, it is reasonable that photoresist property prediction is limited to using unique empirical models developed for each photoresist and device.

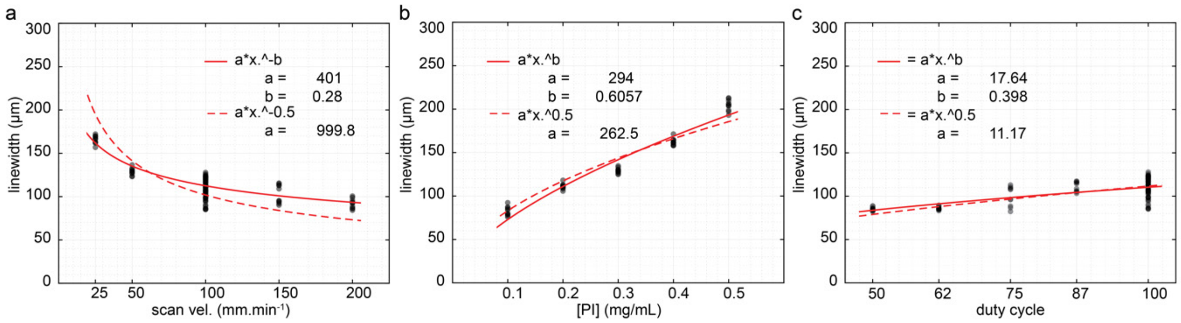

**Fig. S 7 |** The analysis of the FL variables of scan velocity, photoinitiator concentration, and duty-cycle on the resulting polymerized linewidth. The relationship between variables is explored in the context of the standard model of photopolymerization simplified in equation 5, for which the variables should approximate a ½ power relationship with linewidth. The deviations from ½ order show that simplifications can not be applied (sufficiently low *[PI]* and *D* is not met) with additional potential complexities, including that from inhibitory and diffusive species. This exemplifies the complexity of dependent variable prediction for photopolymerization.

**Generating niches from interpolated properties.** To demonstrate the utility of this method, we printed niche filament arrays with adjacent filaments of increasing or decreasing relative Young's modulus, linewidth, and relative bioconjugate fluorescence (**Fig. S 8**). While eight possible permutations with parallel and anti-parallel changes to the three dependent variables are possible (**Fig. S 8**), four were fabricated, as the remaining permutations constitute the mirror-images of those shown (i.e., can be printed by simply reflecting print order, rather than the conditions of fabrication). The fabrication conditions are tabulated at the figure base with properties interpolated on the dotted lines-of-best-fit. Accordingly, the deviation from lines-of-best-fit as seen in the experiment (as run in triplicate), reflect the reproducibility of the method and precision for achieving arbitrarily interpolated material properties. These deviations likely represent the complex non-linear behaviors observed in photochemical reactions as discussed above.

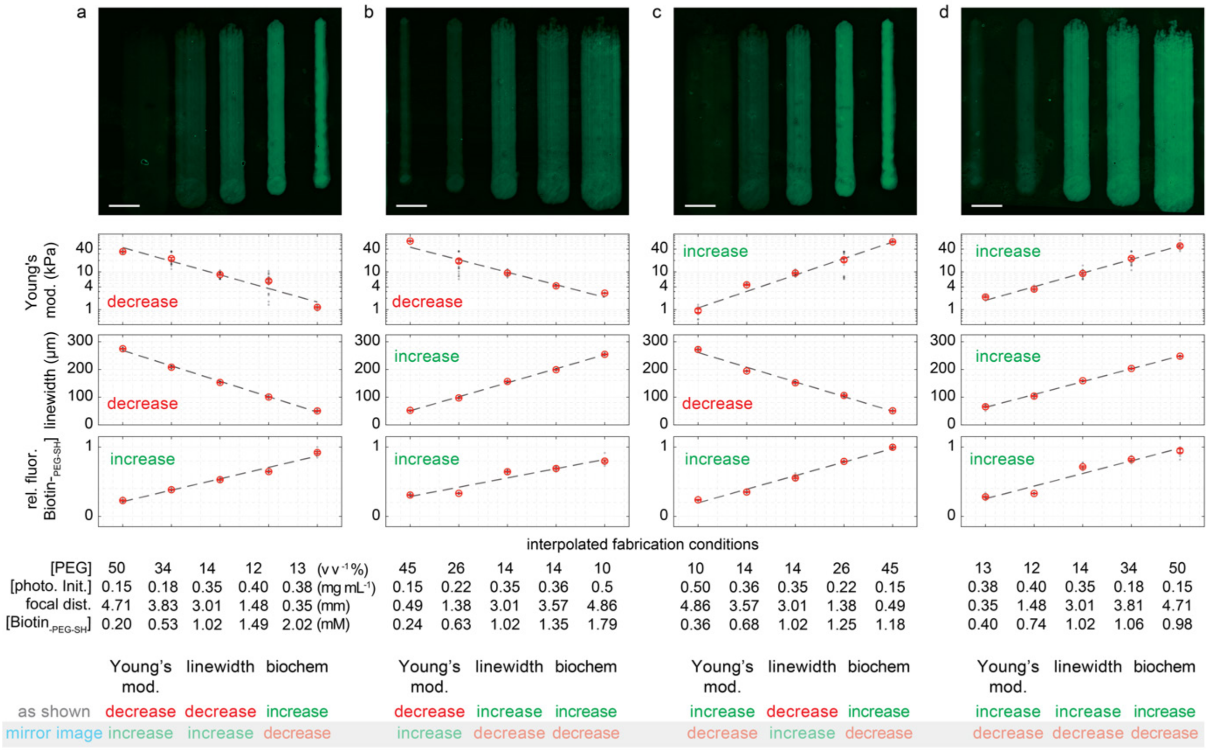

**Fig. S 8 |** Shows microstructured niches generated with interpolated properties from the reduced state-space, with interpolation of the different permutations of the 3 dependent variables either increasing and decreasing in a parallel and antiparallel fashion. At the figure base is the key relating permutation 'mirror images' explicitly, i.e., superposition by reflecting print order.

**Observations of hADSC morphology on [RGD] microstructured niche filaments.** In addition to the characterization of YAP N:C (**Fig 3, Fig. S 9**), the confocal microscopy of human adipose-derived stromal cells (hADSCs) completed over filaments of differing [RGD] allowed the characterization of cell morphological parameters. The below graphs supplement the data provided in **Fig. 3**, showing that structured [RGD] altered a range of morphological parameters, including: the projected cytoplasmic area (**Fig. S 9b**), including cytoplasmic volume (**Fig. S 9c**), nuclear-projected area (**Fig. S 9e**), nuclear volume (**Fig. S 9f**), major axis length 2D (**Fig. S 9g**), cytoplasmic solidity (**Fig. S 9h**), longest axis length 3D (**Fig. S 9i**), nuclear solidity (**Fig. S 9j**). In general, these quantitative morphological characterizations show that increases to [RGD] led to cells with more elongated shapes of larger areas and volume.

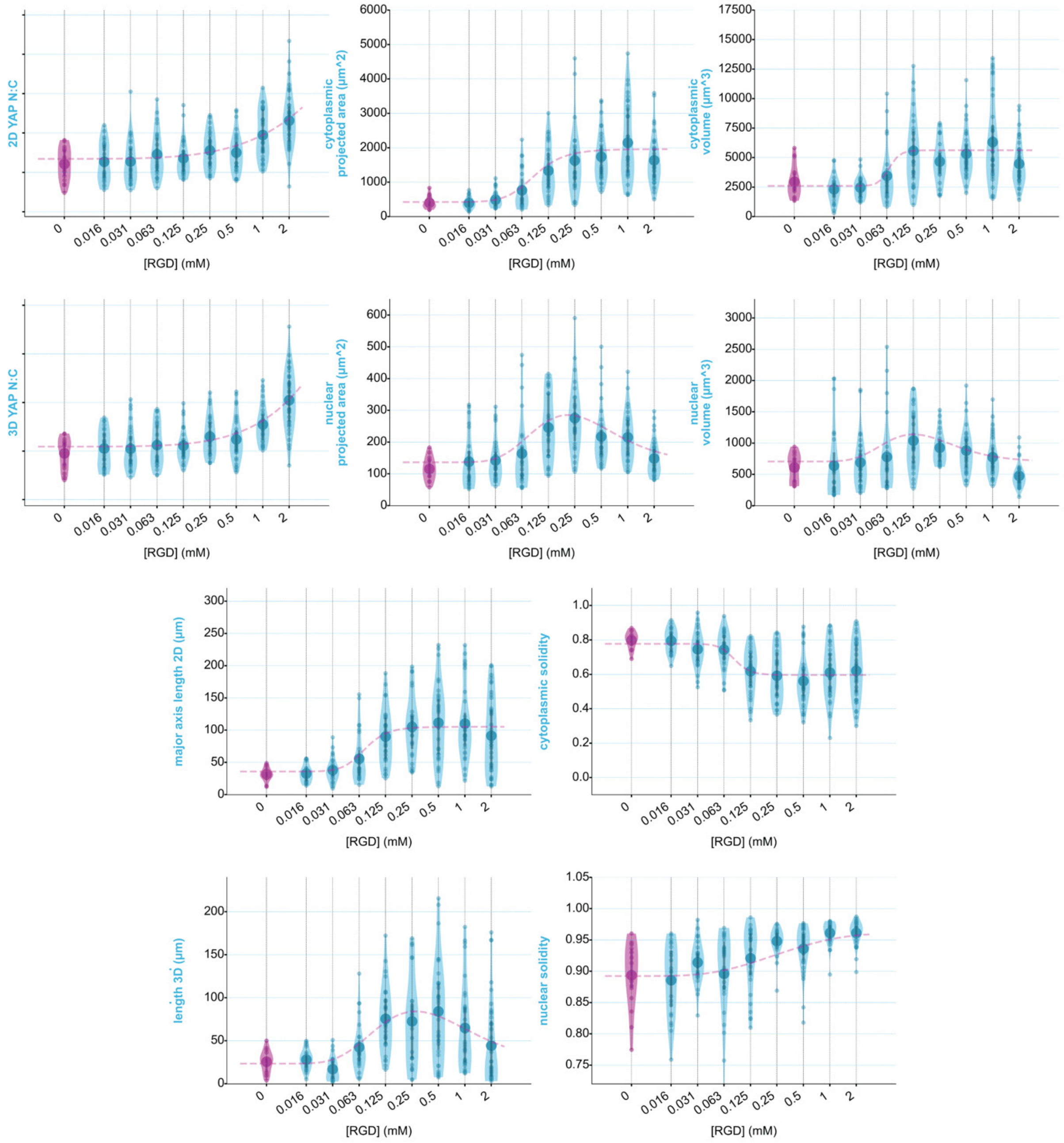

**Fig. S 9 |** Morphological properties of ADSCs as adherent to different [RGD]-structured niche filaments.

**Niche fabrication conditions with defined linewidth, RGD, Young's modulus, and BMP2, as well as bone-fat and germ-layer tissue assemblies**. The below tabulates the interpolated fabrication conditions used to generate the various niche arrays throughout this work.

**Table S 2.** Specific photoresist and MCFL 3DP variables used for fabrication of FL hydrogels with variable Young's modulus, RGD and BMP2 concentrations

| **Label** | **[RGD] (mM)** | **Young's mod. (kPa)** | **[BMP2] (ng/mL)** | **linewidth (µm)** | **PEG ($v\ v^{-1}$ %)** | **[LAP] ($mgmL^{-1}$)** | **focus (mm)** | **biochemical modifier** |
|---|---|---|---|---|---|---|---|---|
| **Fig 1hj** | | | | | | | | |
| 3D structures | (photoabsorber/ Tartrazine = 3 mM) | - | - | 250 | 24.16 | 0.23 | | |
| **Fig 3a** | | | | | | | | |
| 50 µm | 4.4732/0 | 8 | 0 | 50 | 24.99 | 0.22 | 0.5326 | 1.1183 |
| 100 µm | 4.8556/0 | 8 | 0 | 100 | 13.18 | 0.36 | 1.5128 | 1.2139 |
| 150 µm | 4.5364/0 | 8 | 0 | 150 | 13.60 | 0.35 | 3.0025 | 1.1341 |
| 200 µm | 4.1108/0 | 8 | 0 | 200 | 21.85 | 0.25 | 4.3207 | 1.0277 |
| 250 µm | 4/0 | 8 | 0 | 250 | 24.16 | 0.23 | 5.0718 | 1 |
| **Fig 3c, Fig 4c,d** | | | | | | | | |
| stiff-1 | 1.2651 | 20 | 0 | 250 | 35.22 | 0.17 | 4.9879 | 0.84341 |
| stiff-2 | 1.3415 | 15 | 0 | 250 | 31.57 | 0.18 | 5.0295 | 0.89435 |
| stiff-3 | 1.4816 | 10 | 0 | 250 | 25.07 | 0.22 | 5.0552 | 0.98771 |
| stiff-4 | 1.6872 | 5 | 0 | 250 | 15.25 | 0.32 | 4.9684 | 1.1248 |
| stiff-5 | 1.7346 | 2.5 | 0 | 250 | 12.92 | 0.36 | 4.9320 | 1.1564 |
| **Fig 3b, Fig 4a,b** | | | | | | | | |
| rgd-1 | 8 | 8 | 0 | 250 | 24.16 | 0.23 | 5.0718 | 1 |
| rgd-2 | 2 | 8 | 0 | 250 | 24.16 | 0.23 | 5.0718 | 1 |
| rgd-3 | 0.5 | 8 | 0 | 250 | 24.16 | 0.23 | 5.0718 | 1 |
| rgd-4 | 0.125 | 8 | 0 | 250 | 24.16 | 0.23 | 5.0718 | 1 |
| rgd-5 | 0.03125 | 8 | 0 | 250 | 24.16 | 0.23 | 5.0718 | 1 |
| rgd-6 | 0 | 8 | 0 | 250 | 24.16 | 0.23 | 5.0718 | 1 |
| bmp2-1 | 2 | 8 | 0 | 250 | 24.16 | 0.23 | 5.0718 | 1 |
| bmp2-2 | 2 | 8 | 1.6 | 250 | 24.16 | 0.23 | 5.0718 | 1 |
| bmp2-3 | 2 | 8 | 8 | 250 | 24.16 | 0.23 | 5.0718 | 1 |
| bmp2-4 | 2 | 8 | 40 | 250 | 24.16 | 0.23 | 5.0718 | 1 |
| bmp2-5 | 2 | 8 | 200 | 250 | 24.16 | 0.23 | 5.0718 | 1 |
| bmp2-6 | 2 | 8 | 1,000 | 250 | 24.16 | 0.23 | 5.0718 | 1 |
| **Fig 4i,j,k** | | | | | | | | |
| bone µstruc | 3.5774 | 15 | 894.3 | 250 | 31.57 | 0.18 | 5.0295 | 0.8944 |
| fat µstruc | 4.6256 | 2.5 | 0 | 250 | 12.92 | 0.36 | 4.9320 | 1.1564 |
| **Fig 5** | **Matrigel (v/v%)** | | | | | | | |
| dev. org.-1 | 50 | 1 | - | 250 | 10 | 0.25 | 4.98864 | - |
| dev. org.-2 | 50 | 6 | - | 250 | 12.6 | 0.25 | 5.02664 | - |
| dev. org.-3 | 50 | 11 | - | 250 | 15.3 | 0.25 | 5.06464 | - |
| dev. org.-4 | 50 | 30 | - | 250 | 18 | 0.25 | 5.10264 | - |
| dev. org. uniform | 50 | 11 | - | 250 | 15.3 | 0.25 | 5.02664 | - |
| **Fig 6** | | | | | | | | **[BMP4]** |
| dev.org.bmp4-400 | 50 | 11 | - | 250 | 15.3 | 0.25 | 5.02664 | 400 ng/mL |
| dev.org.bmp4-0 | 50 | 11 | - | 250 | 15.3 | 0.25 | 5.02664 | 0 ng/mL |

**Representative images of MCFL bone-fat niches, with channels separated to observe mineralization.** Bone-fat niches were imaged following the multiplexing detection of bone and fat markers. This combined LipidTOX together with the CNA35 Collagen 1A stain, a well-established marker of mature bone[3]. Methods characterizing mineralization were not possible due to an inability to multiplex bone mineralization with Alizarin Red (AZR) staining, together with LipidTOX, due to the shared red fluorescence of LipidTOX and AZR. Below we show 3 representative images of the bone-fat microtissues, with channels LipidTOX (red), DNA/Hoechst (blue), CNA35 (green), and the brightfield images separated (**Fig. S 10)**. Arrows show prominent dark ring regions of the brightfield images where mineralization was observed.

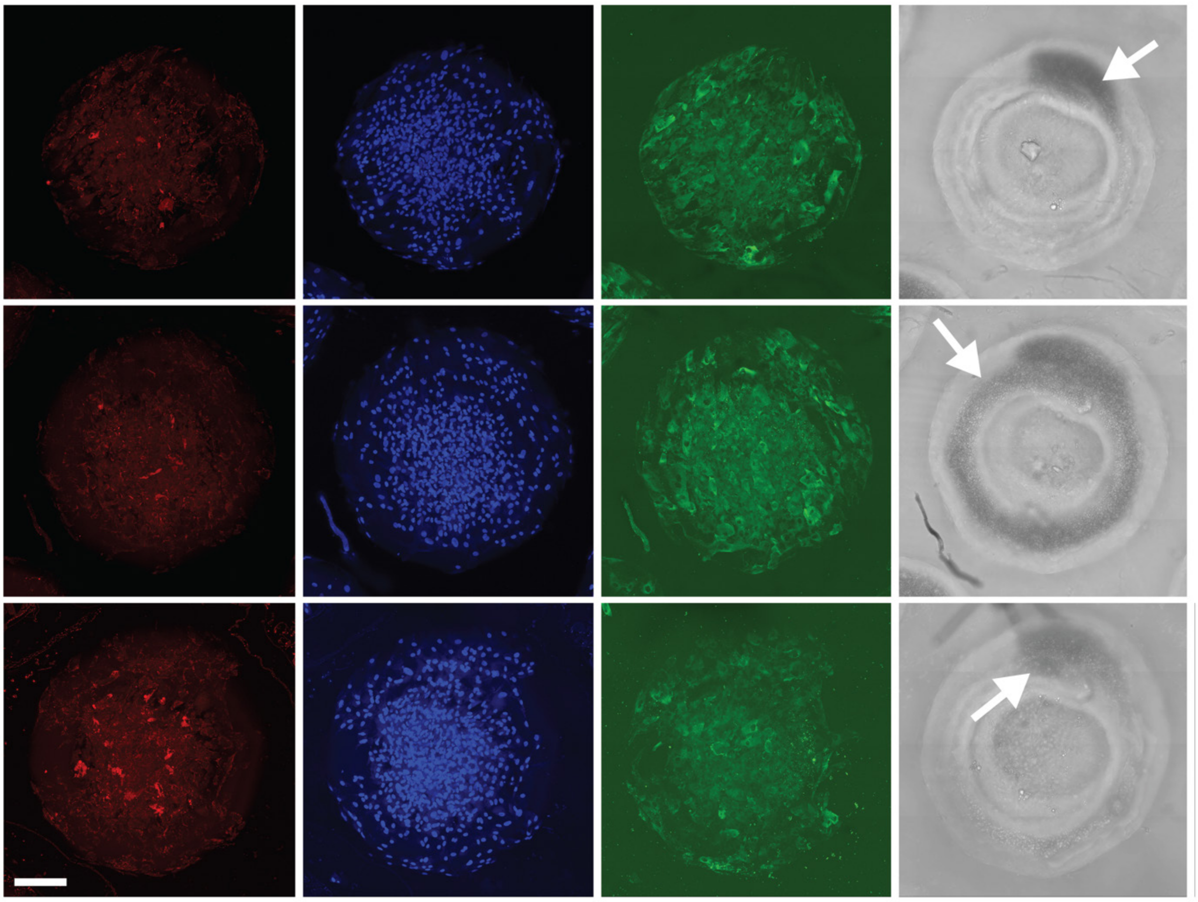

**Fig. S 10 |** Representative images of bone-fat niches with 4 channels separated. Dark regions and arrows indicate region of high mineralization. Gamma corrected for improved visibility of low brightness regions. Scale bar 200 µm.

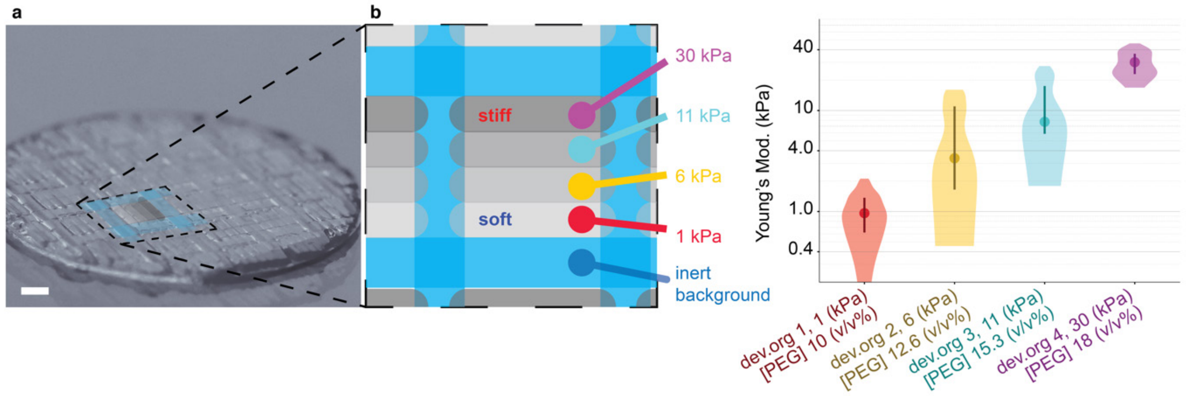

**Fig. S 11 | Force spectroscopic characterization of specific microproperties of mechano-structured niche squares.** Mechano-structured array as fabricated for characterization with AFM. **a,** Macro-lens photography of niche array alongside **b,** schematic of structured mechanical microproperties. **c,** niche properties as measured by AFM force spectroscopy. Scale bar 1 mm.

**Generation of immunofluorescent marker expression maps.** The expression maps in **Fig. 5-6** pool data of marker expression across replicates into a single plot. This was achieved as outlined graphically in the below **Fig. S 12**. First, confocal scans were taken for each marker of interest, as well as the Hoechst nuclear stain, used to generate a mask for each nucleus (**Fig. S 12, step 1**). Using a 3D adaption of the StarDist segmentation tool[4,5], masks for single nuclei were identified (**Fig. S 12, steps 2-3**). Using the nuclei masks, the mean fluorescent intensity for each channel and nuclei was then calculated (**Fig. S 12, step 4**). For the YAP N:C ratio measured in hiPSCs, the cytoplasmic mask was defined by dilating the nuclear volume by half the nucleus volume equivalent radius and subtracting all nuclear masks. Then with the respective mean fluorescent intensity of each marker per nuclear or cytoplasmic masks, the coordinates of each nucleus within the niche are calculated. Linear transformations of each mask's coordinates were mapped to universal coordinates to correct rotational and translational imprecisions when imaging replicates (**Fig. S 12, step 5**). This produced a data set with coordinates and marker intensity for each nucleus (**Fig. S 12, step 6**). Further, experimental replicates and immunostained markers that could not be multiplexed could be pooled, plotted, and compared (**Fig. S 12, step 7**). 3D scatter and surface plots as shown in **Figs 5-6** and **Fig. S12-15, S21, S23** were then generated using matplotlib. Within plots, each dot represents a single cell with the height, size, and transparency of the dot scaled to the normalized fluorescent intensity of the marker across replicates, with further normalization on the total dot area in each plot. Surface plots were then fit to the local mean of the marker expression for the given scatter plot. Additionally, to limit crowding, the number of dots/cells was limited to 200,000, with cells chosen randomly to this limit if exceeded. Further, for merge plots where multiple markers are shown simultaneously, a maximum number of 60,000 cells selected at random are shown for each marker. 3D scatter, and surface plots are shown alongside representative fluorescent microscopy, for which saturation correction has been applied for improved visibility. Specifically, clipping and linear normalization to a maximum saturation of a 98% percentile bit-depth value used for the representative confocal MIP images shown in **Fig 5, Fig. 6,** and all supplemental figures showing hiPSCs.

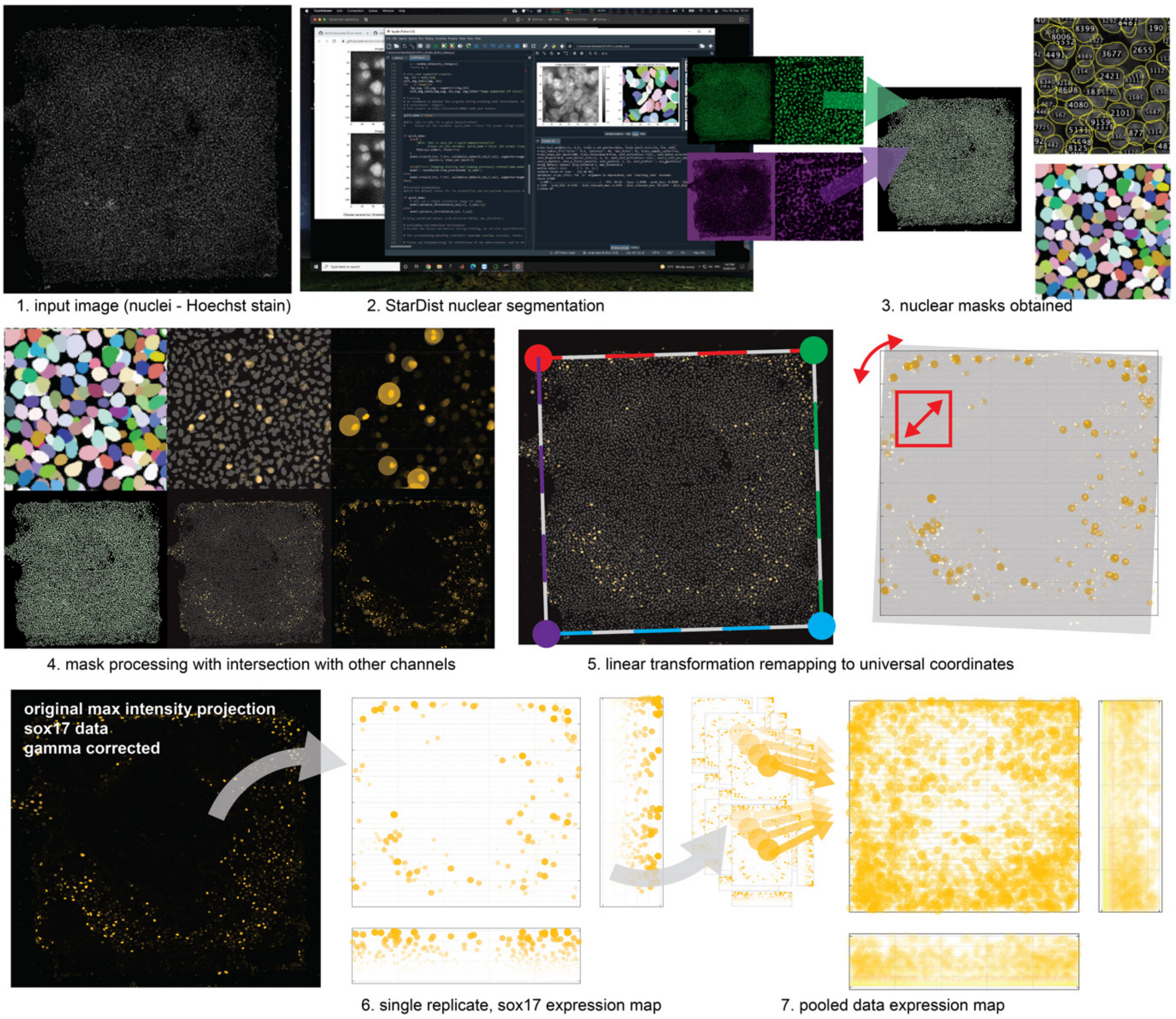

**Fig. S 12 |** Overview of characterization method used to generate tissue expression maps with pooled and mapped data across replicates.

**Replication and comparison of tissue patterning with circular glass-control experiments.** To establish a protocol for using niches with structured microproperties, we first aimed to replicate the seminal work in embryonic micropatterning completed by Warmflash et al. 2015[6]. We first replicated the pattering of germ-layer derivatives in circular tissue micropatterns *in vitro*, with the culture of hiPSCs (HPSI0314i-hoik_1) plated on a Matrigel-coated glass 1000 µm diameter circular adhesive templates, with boundaries defined using inert photo-printed PEG (**Fig. 5a**). Cells were seeded in Essential 8 Flex media, with CEPT cocktail[7] for 2 h as per method indicated on **Fig. 5** and **Fig. 6** respectively. Cells cultured on the homogenous circular template generated radially symmetric germ-layer derivatives (**Fig. S 13**). Measuring tissue-patterning with progenitor cells of the mesoderm (BRA-positive), endoderm (SOX17-positive), and ectoderm (SOX2-positive) at 48 h revealed a phenotype equivalent to the Warmflash work's time point at 24-30 h. Accordingly, the endpoint of our protocol was extended from 48 h to 72 h. **Fig. S 13** below compares 48 h and 72 h time points, showing representative images, alongside the pooled-position-mapped marker expression. At 48h, expression of SOX17 was only found at the periphery, with maximum intensity levels comparable to background staining. Further, BRA expression was only apparent in the most peripheral cells. The extension of our endpoint to 72 h produced BRA expression became relatively centralized, and SOX17 expression became prominent from the background levels. In subsequent experiments exploring the use of niche mechano-chemical properties for directing cell fate decisions, the endpoint was extended to 72 h, except for **Fig. S 15** below, showing tissue patterning marker expression in mechano-structured niches at 48 h.

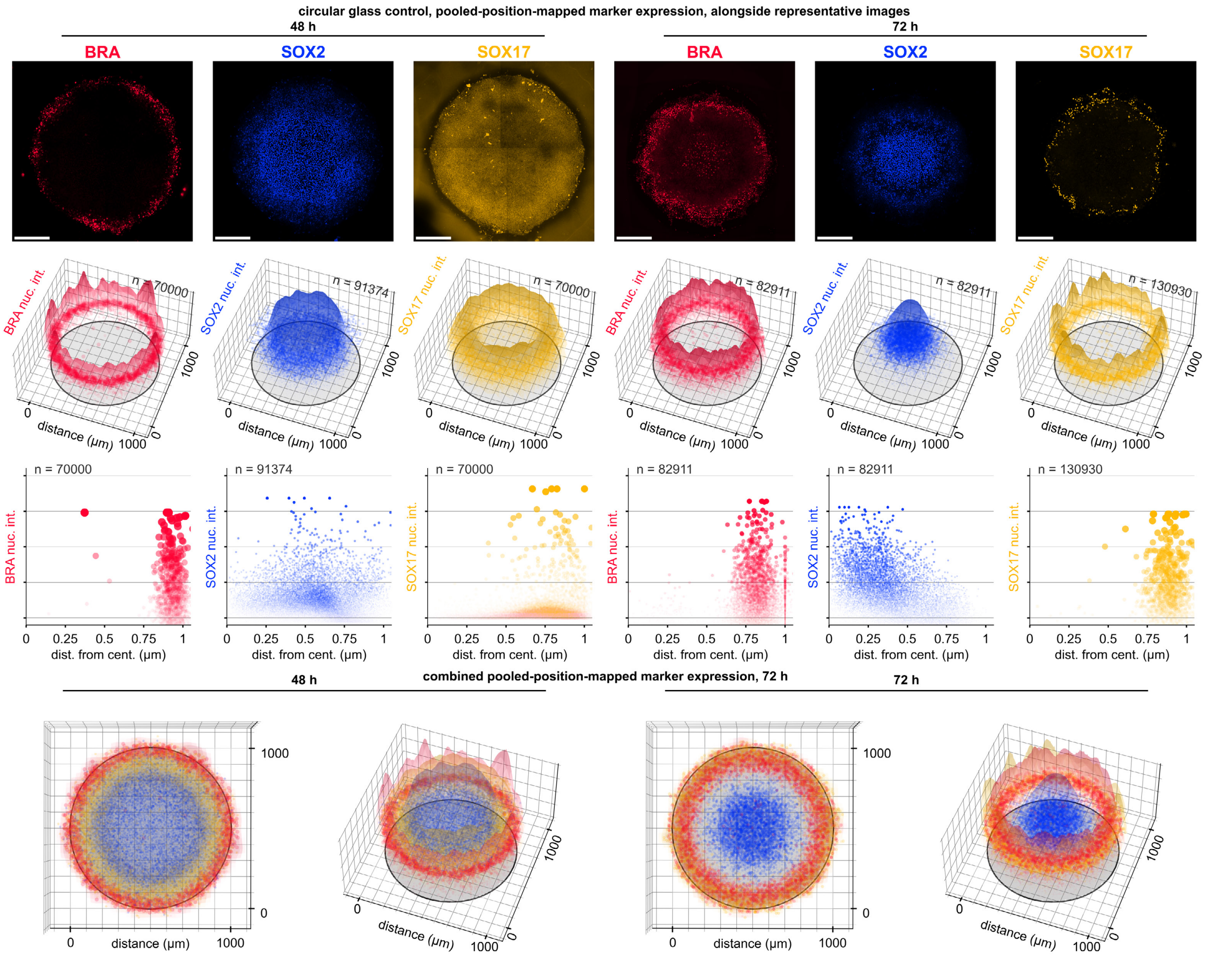

**Fig. S 13 | Comparison of hiPSC culture endpoint by replicating circular glass-control experiments. Row 1** shows representative images of circular templated germ-layer tissue patterning with Matrigel-coated-glass and inert printed template boundaries. Immunofluorescent staining of templates 48 h (three panels at the left) and 72 h (three panels at the right) post-BMP4-treatment (50 ng/mL), showing (mesoderm: BRA, red; ectoderm: SOX2, blue and endoderm: SOX17, yellow). For improved visibility, images are shown with linear-scaling and clipping to a maximum saturation of a 98% percentile bit-depth value. Scale bar 200 µm. **Row 2** shows position-mapped expression graphs of replicate pooled-data for respective markers labeled directly above, with dot color and size scaled as described in the supplementary information above. **Row 3** shows the marker expression intensity projected along the radial dimension of the circular pattern. Each dot represents a single cell with color and size scaled as described in the supplementary information above. **Row 4** shows two views of tissue patterning at both 48 h (two panels at the left) and 72 h (two panels at the right), as shown in *Row 2*, with corresponding markers overlaid on a single plot. Notably, the 72 h endpoint shows similar tissue patterning to Warmflash et al. obtained at 48h.

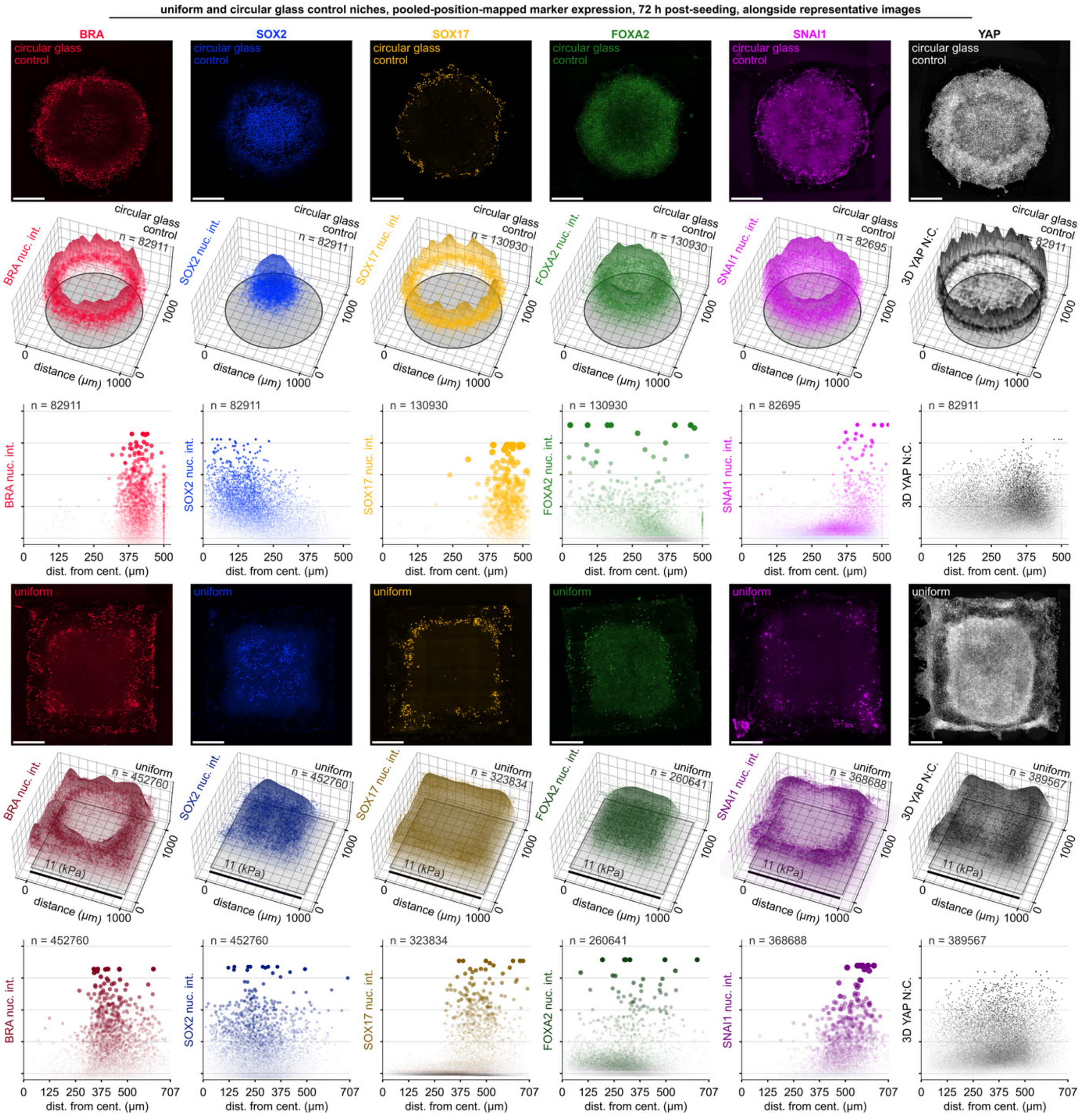

**Fig. S 14 | Comparison of circular glass control niches with square uniform niches at 72 h. Rows 1 and 4** show representative images over circular glass and uniform niches at 72 h for the markers: BRA-red, SOX2-blue, SOX17-yellow, FOXA2-green, SNAI1-purple, and YAP-grey. The overlaid text indicates the underlying niche type. **Rows 2 and 5,** Show replicate data presented as 3D scatter and surface plots overlaid on a representation of the corresponding niche. **Rows 3 and 6,** Shows plots with reduced dimensionality, projecting the mapped x/y position into the single dimension - distance from the center of the niche. Distance from the center of the niche is plotted against the relative fluorescence intensity of the given marker. The horizontal axis for circular plots extends to 500 μm in row 3, extending to 707 μm for square plots (the maximum distance possible from the center of a square with 1000 μm edges). Patterning over the circular glass and uniform niches are similar, with peripheral mesodermal and endodermal markers.

In addition to the characterization of a range of marker expression levels over niches with uniform or mechano-structured properties at an endpoint of 72 h, we stained and characterized niches at 48 h. Mechano-structured niches were fabricated with a 1D gradient in their mechanics as per **Fig 5** of the main article (**Fig. 5d**) alongside a control niche with uniform microproperties (**Fig. 5a**). hiPSCs cultured on both niche types were induced to differentiate with BMP4 (50 ng/mL) into progenitor cells of the mesoderm (BRA-positive), endoderm (SOX17-positive), and ectoderm (SOX2-positive)[6]. Image analysis pooling replicate-data (**Fig. S 12**) revealed that hiPSCs cultured on uniform stiffness substrate displayed regionalization of germ layer tissues from their center-to-periphery (**Fig. 5c,j,i,k**), similar to that observed for circular micropatterns (**Fig. S 13**)[6]. In contrast, cells cultured over mechano-structured niches were able to reconfigure the tissue pattern. Expression of BRA- and SOX17-positive cells were localized to niche regions of low stiffness and SOX2-positive cells to regions with high stiffness (**Fig. 5f,h,l**). This observation is consistent with the patterning observed at 72 h.

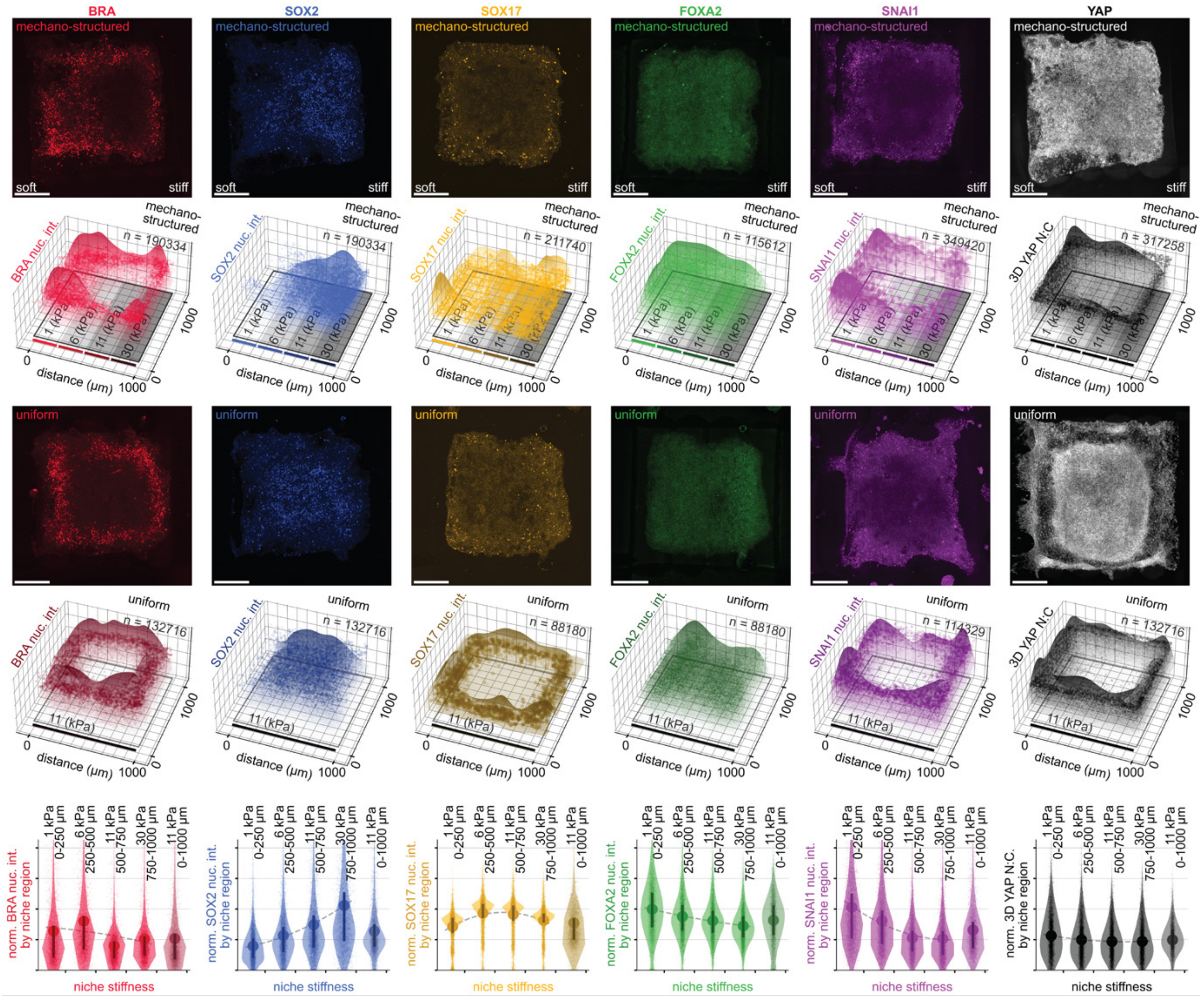

**Fig. S 15 | Niche-mechanics directed marker expression at 48 h. Rows 1 and 3** shows representative images over mechano-structured and uniform niches at 48 h for the markers: BRA-red, SOX2-blue, SOX17-yellow, FOXA2-green, SNAI1-purple, and YAP-grey. Overlaid text indicates image orientation concerning the niche type. The color-coding of markers is consistent throughout this work, with saturation correction applied to representative images, with clipping and linear normalization to a maximum saturation of a 98% percentile bit-depth value applied for improved visibility. Image overlay labels the mechanical properties over the three different niches. The orientation of niches is consistent across the figure, with the 1D mechanical gradient shown soft-to-stiff and left-to-right. Scale bars 200 µm. **Rows 2 and 4,** Show replicate data presented as 3D scatter and surface plots overlaid on a representation of the corresponding niche. Surface plots are fit to the local mean of the marker expression and plotted on a separate axis to the scatter plots. Each dot represents a single cell with the size and transparency of the dot scaled to the normalized fluorescent intensity of the marker across replicates. **Row 5,** Shows violin plots that compare the mean and distribution difference, including mean and 1$^{st}$/3$^{rd}$ quartile lines of replicate data as mapped to respective niche regions indicated on the x-axis.

**Fig. S 16** below shows brightfield imaging of the four different niches used in this work with hiPSCS. Text overlays describe the niche properties, which are consistent between rows. **Fig. S 16 Column 1** shows the circular glass control micropatterns. Following the seeding step, leaving cells to attach for 2 h with the pharmacological cocktail CEPT, hiPSCs at the center of the colony exhibit tight junctions characteristic of hiPSCs. At 72 h, a circular ring structure of the cells appears along the radius of the structure - consistent with the appearance of cells in predicate work[6]. **Fig. S 16**, **Column 2** shows brightfield imaging of cells attached to morphogen-structured niches. Relative to other niches, the lower seeding density used for morphogen- structured niches (450 k cells per $cm^2$) is evident. The 72 h time point shows the formation of circular node-like structures occupied by large cells over BMP4 containing regions, with more densely packed and smaller cells over regions absent of BMP4. **Fig. S 16, Column 3** shows brightfield microscopy of mechano-structured niche structures. The boundary of the niche pattern exhibits larger, less densely packed cells which bias towards the softer regions of the substrate. Compared to the soft regions on the left, the stiff boundary at the right exhibits relatively small cells with dense packing. When compared with the uniform niche, cell morphology and density are symmetric around the square structure.

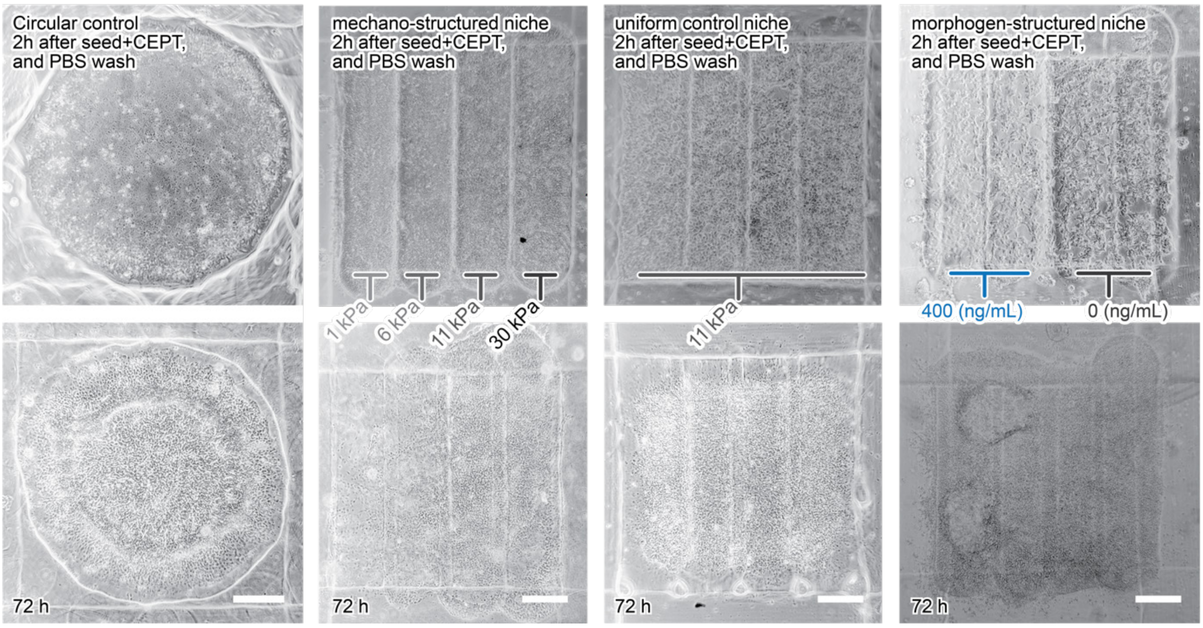

**Fig. S 16 | Brightfield imaging of the four different niches explored with hiPSCs at 2 and 72 h. Row 1** Shows representative images of niches at 2 h, with text overlays describing the niche properties that are consistent between rows. **Row 2** shows niches at 72 h. **Column 1** shows circular glass control experiments. **Column 2** mechano-structured niches. **Column 3** shows uniform niches. **Column 4** shows morphogen-(bmp4)-structured niches.

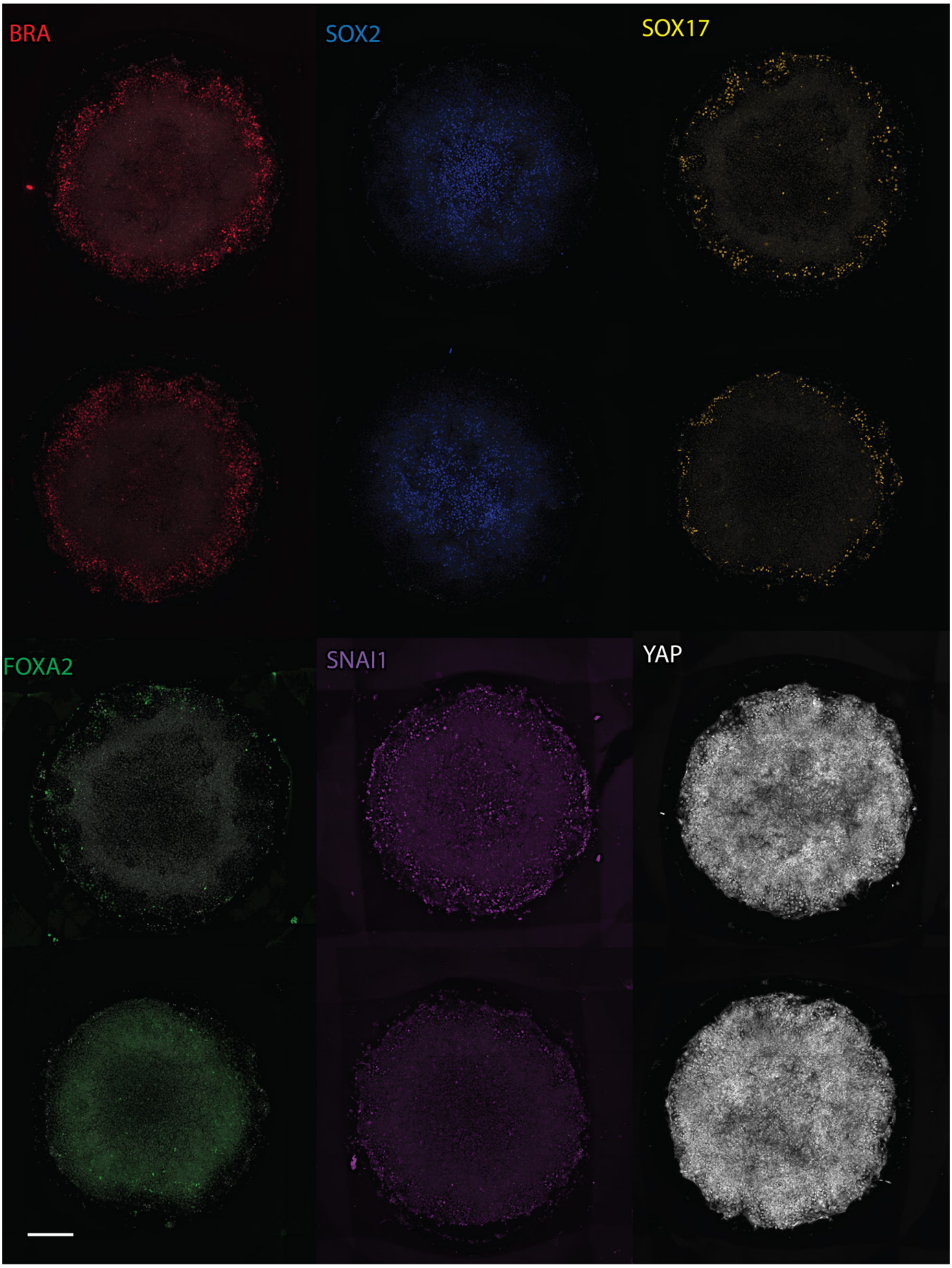

410 **Fig. S 17 | High resolution images of representative circular coverslip control niches at 72 h.** Shows high larger/high magnification images of markers at 72 h, overlaid on gray nuclear stain. Scale bar 200 µm. The color-coding is consistent though work. Saturation correction applied to above images, with clipping and linear normalization to a 99.8% percentile bit-depth for improved visibility.

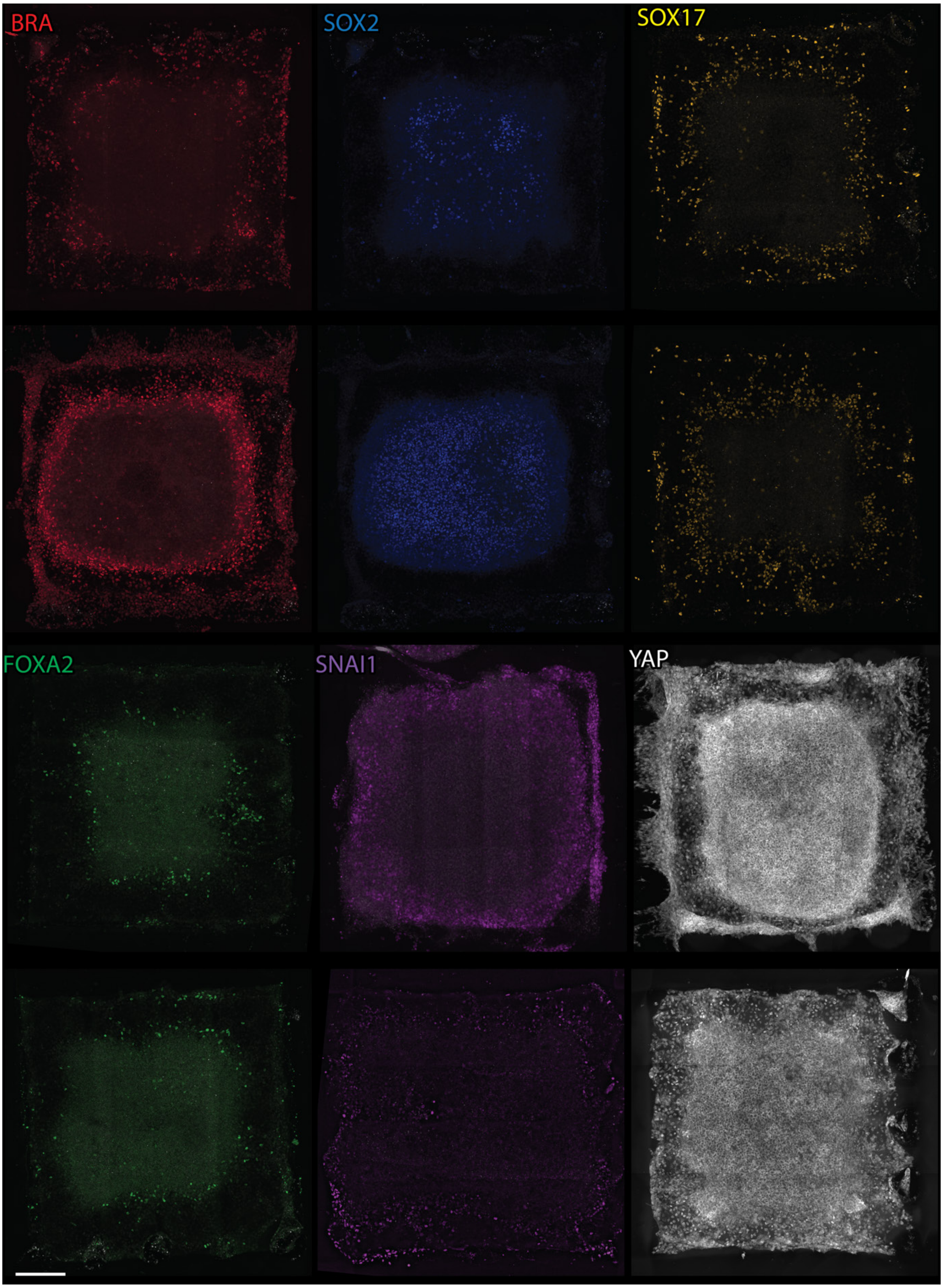

415 **Fig. S 18 | High resolution images of representative uniform control niches at 72 h.** Shows high larger/high magnification images of markers at 72 h, overlaid on gray nuclear stain. Scale bar 200 µm. The color-coding is consistent though work. Saturation correction applied to above images, with clipping and linear normalization to a 99.8% percentile bit-depth for improved visibility.

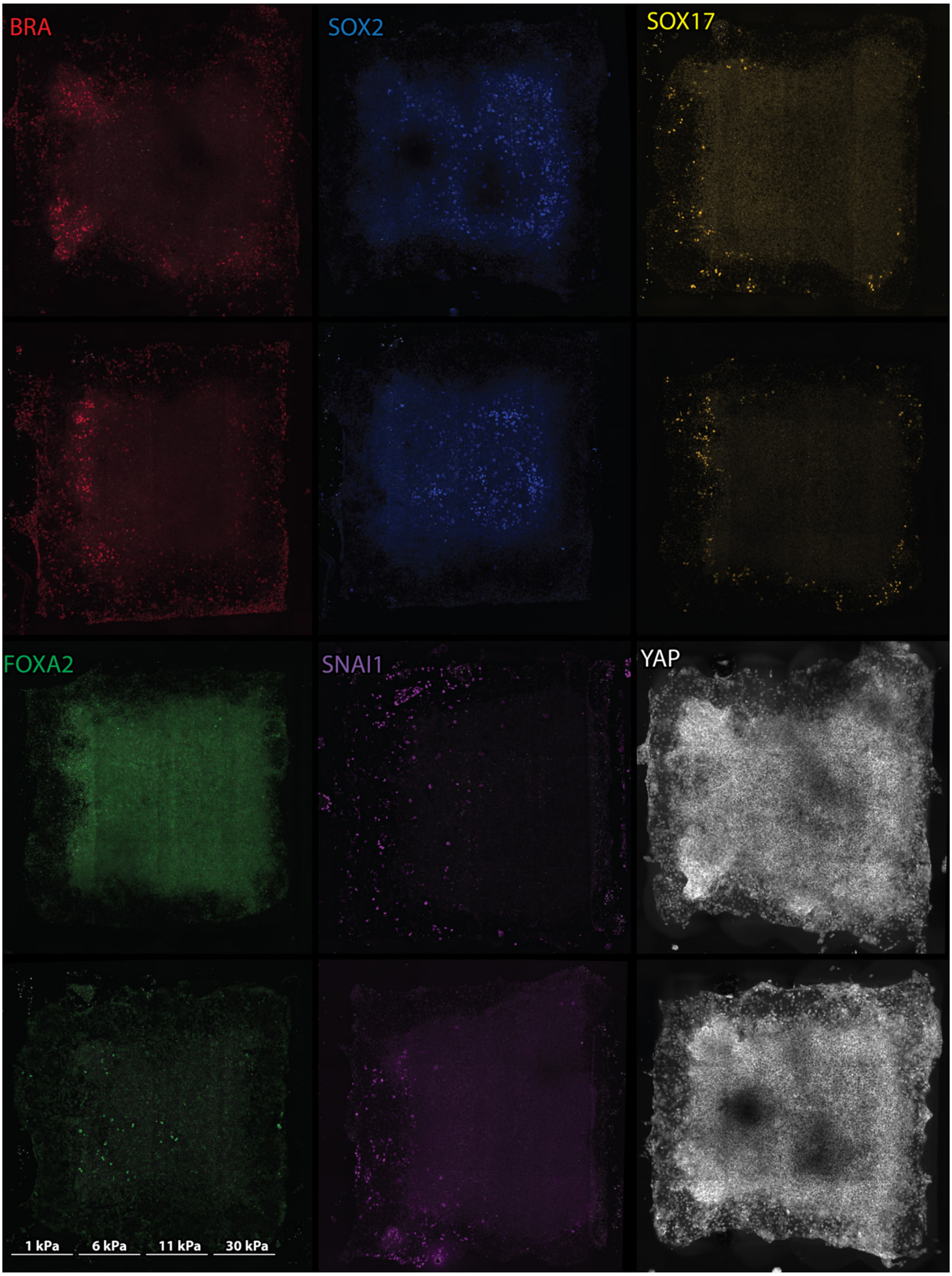

**Fig. S 19 | High resolution images of representative mechano-structured niches at 72 h.** Shows high larger/high magnification images of markers at 72 h, overlaid on gray nuclear stain. Niches are printed at 1000 μm across. The color-coding is consistent though work. Saturation correction applied to above images, with clipping and linear normalization to a 99.8% percentile bit-depth for improved visibility.

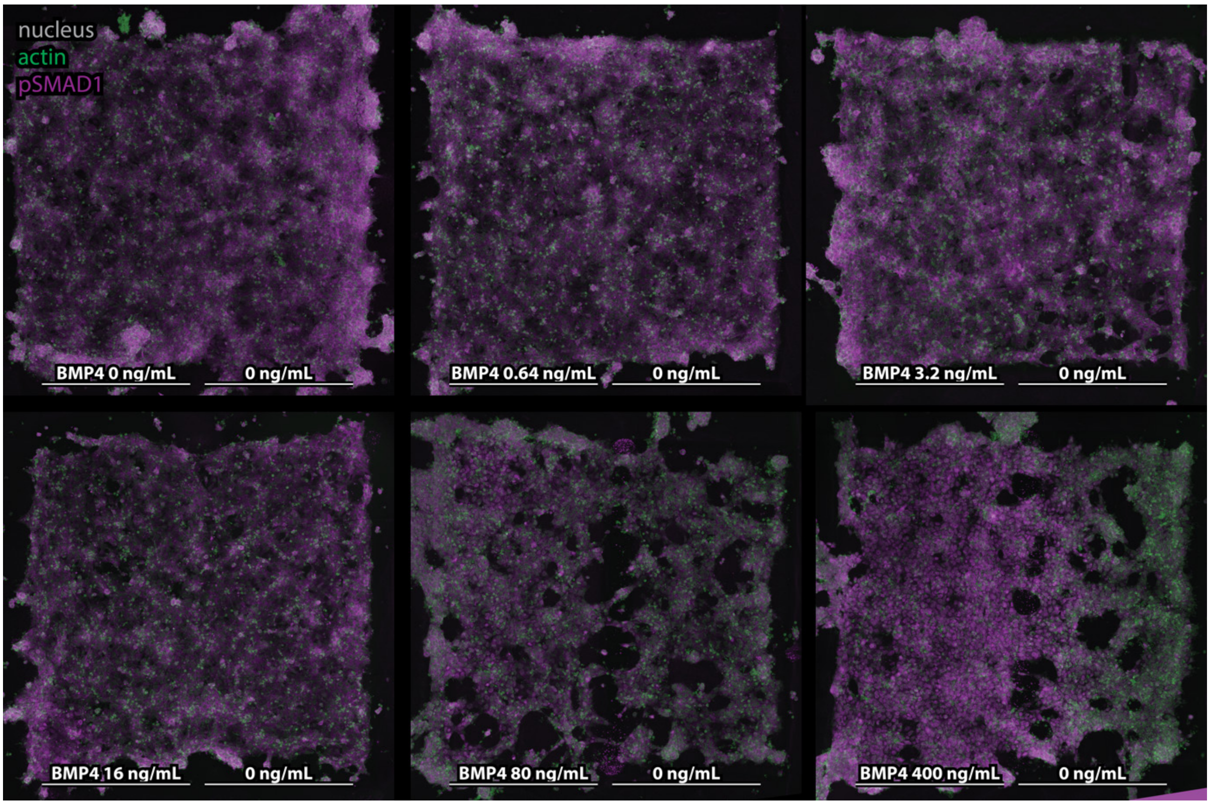

**Fig. S 20 | High resolution images of representative morphogen-structured niches at 2 h.** Shows high larger/high magnification images of pSMAD1 at 2 h, overlaid with gray nuclear stain, and green actin stain. Niches are printed as 1000 µm across. Saturation correction applied to above images, with clipping and linear normalization to a 99.8% percentile bit-depth for improved visibility.

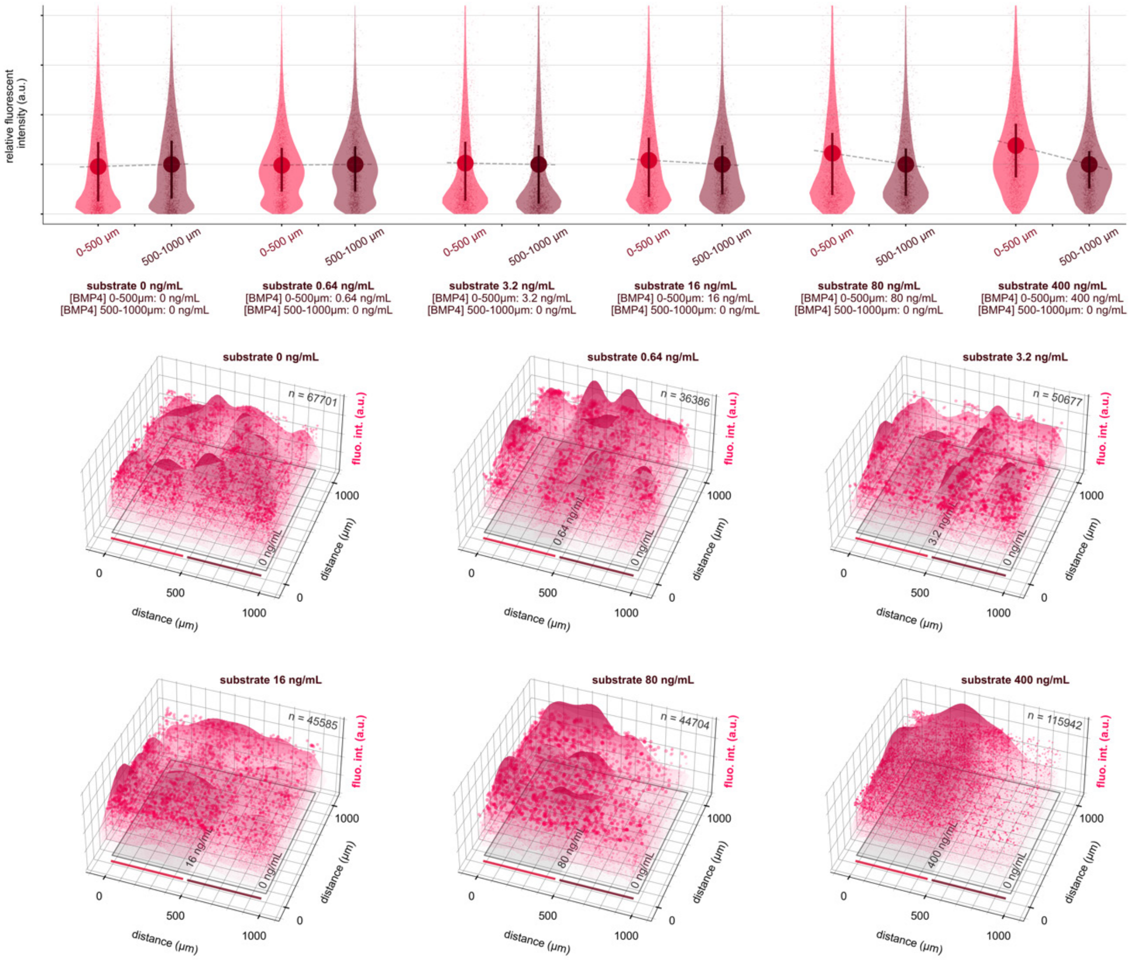

**Fig. S 21 | Comparison of pSMAD1 N:C ratio for varying concentrations of BMP4.** Violin plots compare the mean and distribution differences of pSMAD1 N:C ratio for different concentrations of BMP4 added to the morphogen-structured half of the niche at 2 h post-seeding, as normalized to the N:C ratio over the niche half lacking BMP4. Substrate concentrations of BMP4 are shown for violin plot pairs along the horizontal axis. 3D scatter, and surface plots are shown for replicate-pooled data, with titles showing substrate concentration of BMP4, overlaid on a representation of the corresponding niche properties. A dose-dependent increase to the relative N:C ratio over regions with BMP4 was observed with increasing [BMP4].

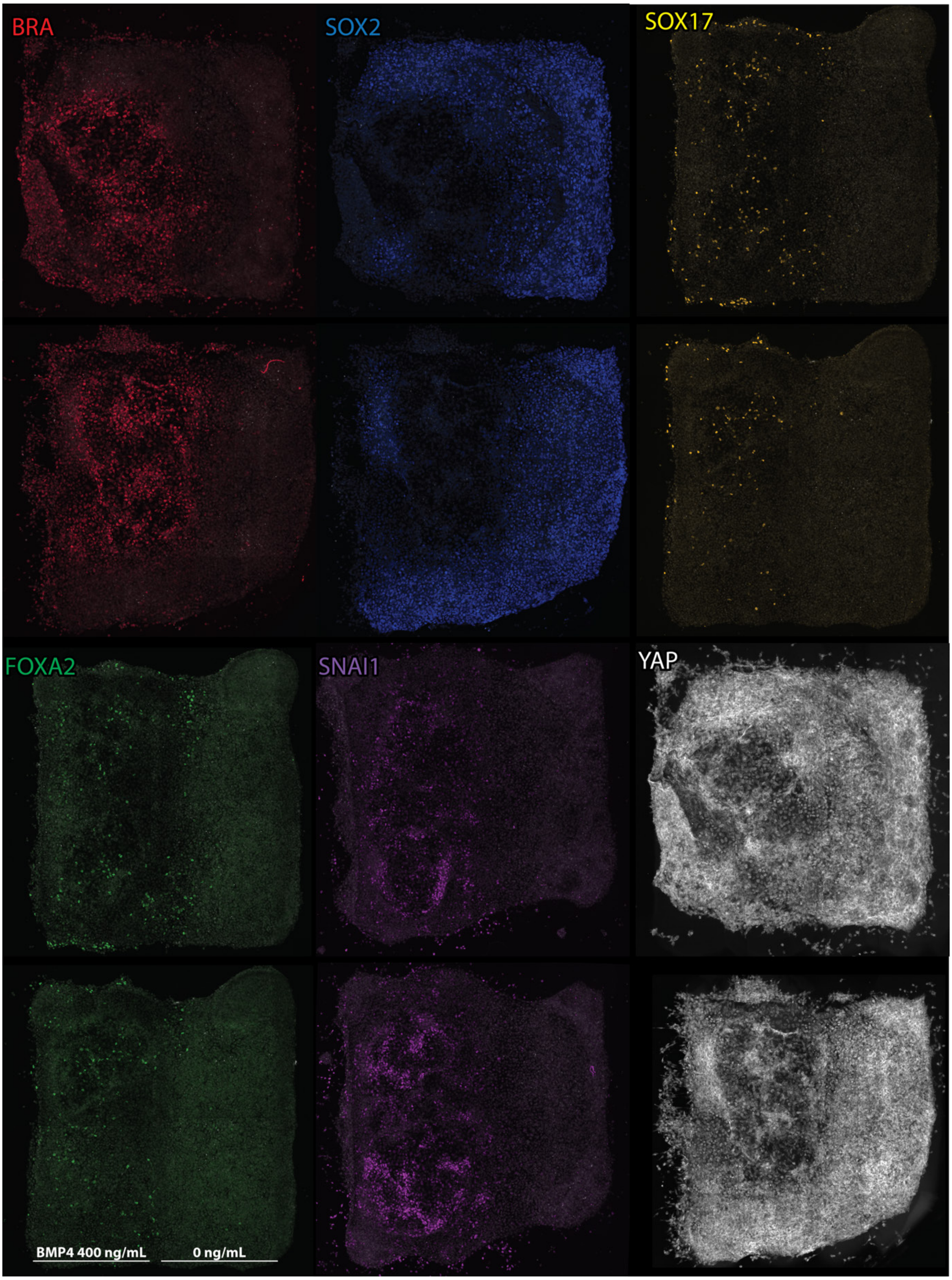

**Fig. S 22 | Representative morphogen-structured niches at 72 h.** Shows high larger/high magnification images of markers at 72 h, overlaid on gray nuclear stain. Niches are printed as 1000 µm across. Scale bar 200 µm. The color-coding is consistent though work. Saturation correction applied to above images, with clipping and linear normalization to a 99.8% percentile bit-depth for improved visibility.

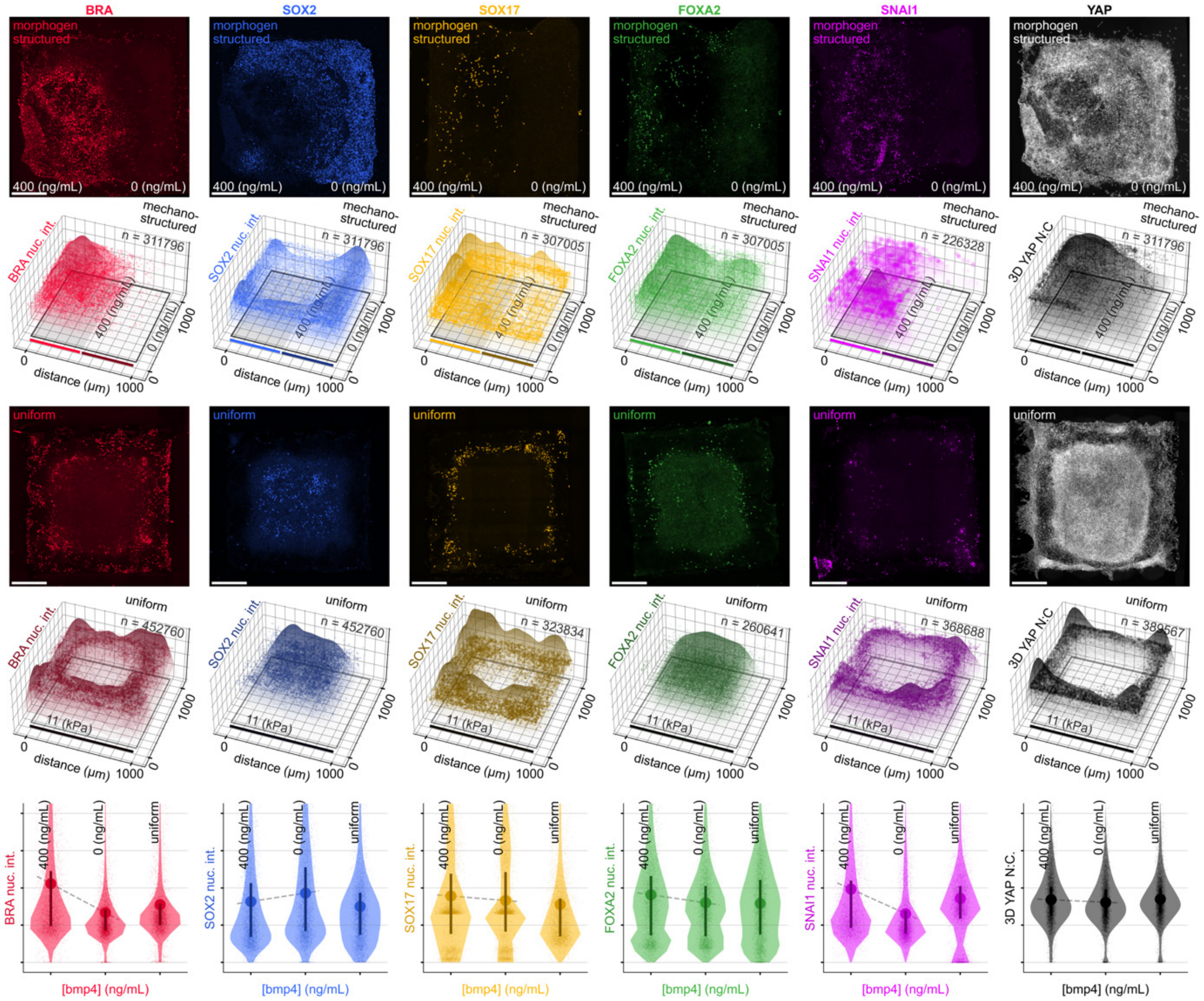

**Fig. S 23 | Comparison of morphogen-structured and uniform niches at 72 h. Rows 1 and 3** shows representative images with overlaid text indicating the underlying niche type and niche orientation. The orientation of niches is consistent across the figure, showing BMP4 containing regions at figure left and regions absent of BMP4 at the right. **Rows 2 and 4,** Show replicate data presented as 3D scatter and surface plots overlaid on a representation of the corresponding niche. **Row 5,** Violin plots compare the mean and distribution difference, including mean and 1st/3rd quartile lines of replicate data mapped to respective niche regions.

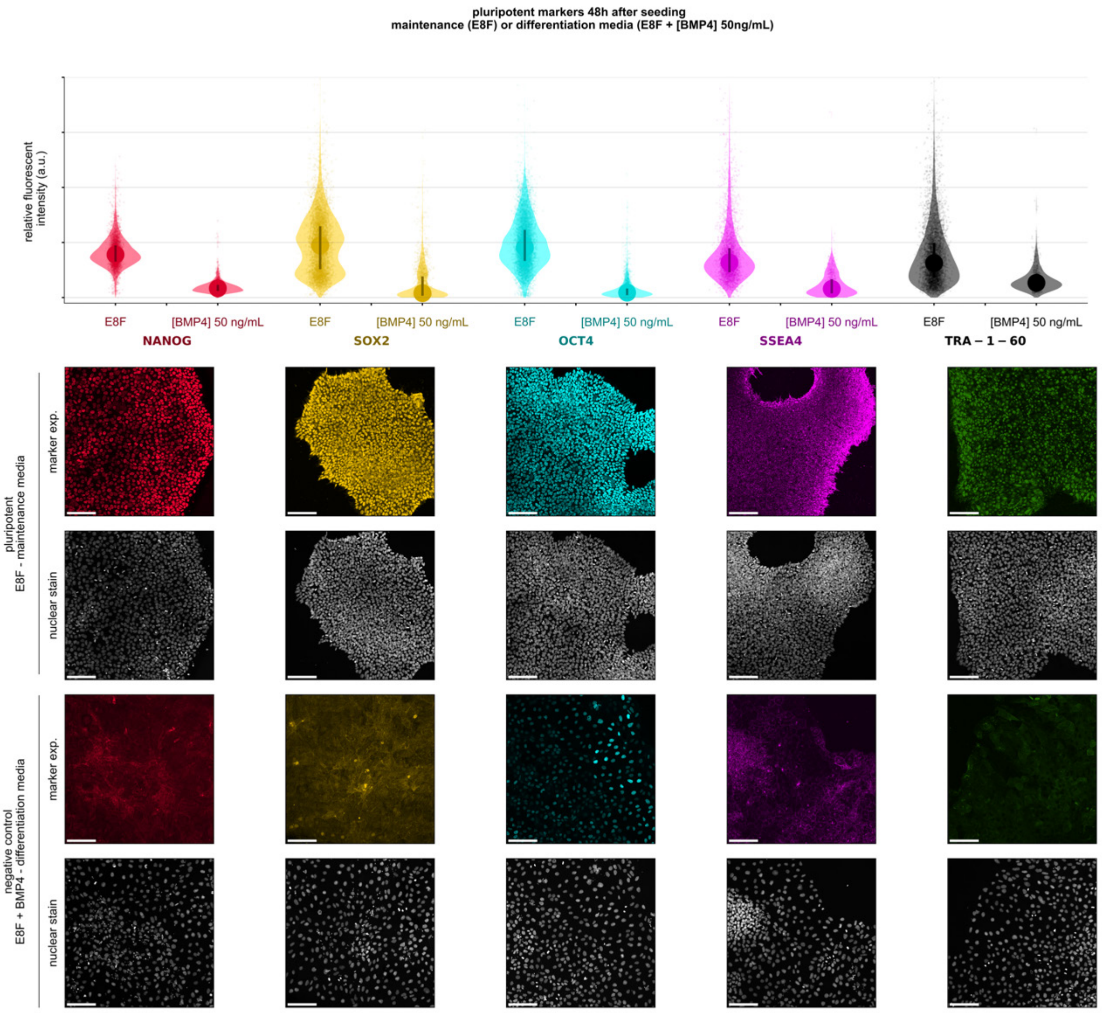

**Fig. S 24 | Confirmation of hiPSC pluripotency as compared with differentiated negative control.** Shows quantification and representative images of pluripotent markers confirmed upon first receiving the HOIK1 cell line in passaged cell colonies comparing Essential 8 Flex (E8F) media or E8F media with BMP 4 (50 ng/mL).

### References

1. Grigoryan, B. *et al.* Multivascular networks and functional intravascular topologies within biocompatible hydrogels. *Science* **364**, 458–464 (2019).

2. Weiss, P. Principles of polymerization, George Odian, Wiley-Interscience, New York, 1981, 731 pp. *J. Polym. Sci. Polym. Lett. Ed.* **19**, 519–519 (1981).

3. Chan, C. K. *et al.* Identification of the human skeletal stem cell. *Cell* **175**, 43–56 (2018).

4. Schmidt, U., Weigert, M., Broaddus, C. & Myers, G. Cell detection with star-convex polygons. in 265–273 (Springer, 2018).

5. Weigert, M., Schmidt, U., Haase, R., Sugawara, K. & Myers, G. Star-convex polyhedra for 3d object detection and segmentation in microscopy. in 3666–3673 (2020).
6. Warmflash, A., Sorre, B., Etoc, F., Siggia, E. D. & Brivanlou, A. H. A method to recapitulate early embryonic spatial patterning in human embryonic stem cells. *Nat. Methods* **11**, 847 (2014).
7. Chen, Y. *et al.* A versatile polypharmacology platform promotes cytoprotection and viability of human pluripotent and differentiated cells. *Nat. Methods* **18**, 528–541 (2021).
